# MagPiezo: a Magnetogenetic Platform for Remote Activation of Endogenous Piezo1 Channels in Endothelial Cells

**DOI:** 10.1101/2025.06.30.661957

**Authors:** Susel Del Sol-Fernández, Mariarosaria De Simone, Yilian Fernández-Afonso, Daniel Garcia-Gonzalez, Pablo Martínez-Vicente, Thomas S van Zanten, Raluca M. Fratila, Maria Moros

## Abstract

This work introduces MagPiezo, a novel magnetogenetic platform designed to enable remote and targeted manipulation of endogenous Piezo1 channels without genetic modification. MagPiezo integrates a custom-built magnetic applicator and 19 nm octahedral mixed ferrite nanoparticles (MFs) with enhanced magnetocrystalline anisotropy, engineered to selectively target Piezo1 channels. The application of a low intensity (< 40 mT), low frequency (1 Hz) magnetic field is sufficient to generate wireless torque forces in the piconewton range capable of gating Piezo1 channels in vitro. Remarkably, applying MagPiezo for short times (< 1 h) induces robust responses in endothelial cells, including calcium influx, cytoskeletal reorganization and transcriptional activation. Unlike other magnetogenetic tools, MagPiezo avoids the need to assemble the magnetic nanoparticles (MNPs) into bigger beads to reach the opening threshold and does not need to genetically engineer Piezo1 channels. Moreover, it uses low magnetic field intensities and frequencies, enhancing its potential for in vivo translation. Therefore, MagPiezo is a powerful platform for manipulating Piezo1 channels and dissecting mechanotransduction events in vascular biology, paving the way for new strategies to modulate mechanoreceptors both in vitro and in vivo.

## 1. Introduction

In a process called mechanotransduction, our cells are able to sense mechanical forces from their environment and convert them into biochemical signals, leading to important cellular processes. Among the key players, Piezos (Piezo1 and 2) have emerged as crucial mechanosensitive (MS) ion channels, highly involved in both physiological and pathological contexts.[1–4] These channels are non-selective cation channels that can open in response to stimuli such as pressure or shear stress, allowing cell depolarization and Ca^2+^ influx. Piezo1 is predominantly found in non-sensory cells, including red blood cells, whereas Piezo2 is highly expressed in mechanosensory cells, such as dorsal root ganglia sensory neurons.[4] In vertebrates, expression of Piezo channels is essential for survival, as they are involved in vital processes such as vessel development, touch, nociception and blood pressure regulation.[3,5] Moreover, they are also involved in other processes like cancer progression or immune cell activation.[6]

To date, significant progress has been achieved in understanding the mechanism of Piezo activation, although the whole mechanism remains elusive.[7,8] Due to their role in many important processes, the selective manipulation of Piezo channels holds great potential as novel therapeutic targets. While much attention has been focused on their role in neuromodulation, another major target could be the cardiovascular system as Piezo channels are implied in cardiovascular development and maintenance,[9] but also contribute to related pathogenesis including atherosclerosis, hypertension, and ischemic injury.[10,11] In this context, promoting angiogenesis and neovessel formation by manipulating Ca^2+^ influx in endothelial cells has been proposed as a novel strategy for therapeutic angiogenesis.[12] As Piezo1 is highly expressed in endothelial cells and its gating mediates Ca^2+^ entrance, the selective manipulation of this receptor could be a promising strategy for the treatment of ischemic disease.

However, despite the significant advances made in the last years and the number of techniques available to tackle mechanoreceptors, activating Piezo1 channels and controlling downstream signaling in vivo remains challenging. For instance, some macroscopic methods such as microaspiration or shear stress, lack the possibility to activate the receptors at a molecular level,[13] while other powerful techniques such as electrophysiology or atomic force microscopy (AFM) cannot be applied in vivo. On the other hand, the limited number of chemical agonists against Piezos and their lack of spatiotemporal control hampers their application in vivo due to possible systemic toxicity. Therefore, new tools for remote control of these MS channels with spatiotemporal resolution are needed.

A powerful approach for manipulating mechanoreceptors to activate downstream cellular processes involves the generation of mechanical forces through the combination of magnetic fields and magnetic actuators such as magnetic particles, an approach coined as magnetogenetics.[14] Similar to optogenetics, that uses light to control cellular function, magnetogenetics is a non-invasive approach. However, unlike light, magnetic fields exhibit high tissue penetrability, which is crucial for the development of untethered deep tissue in vivo applications. [15] For many years, magnetic tweezers have used magnetic microparticles to study how mechanical stimuli influence biological processes, generally at a single-cell level. When compared to magnetic microparticles, smaller MNPs offer advantages, as they show better spatial control at the molecular level.[16] In the context of mechanotransduction, MNPs enable the manipulation of MS channels by applying mechanical forces through two main mechanisms: i) torque generated by changes in the magnetization state, or ii) translational forces induced by the magnetic field gradient. While recent studies have demonstrated the potential of this strategy to mechanically activate mechanoreceptors, the use of ferritin proteins or small MNPs might be compromised by the lower forces that can be generated when compared with larger particles.[17–24] Thus, a whole optimization of the system (MNPs and magnets) is needed.

In the context of magnetogenetics, recent pioneering studies have demonstrated the possibility of manipulating Piezo1 channels for neuromodulation both in vitro and in vivo using magnetic particles and a circular array of magnets.[25–27] However, to generate enough force these studies required the assembly of MNPs on bigger substrates (200-500 nm), involving additional steps. Moreover, in all these studies Piezo1 channels were exogenously expressed in cells, which does not represent reality and likely hampers their further clinical translation.[25–27] Additional hurdles for targeting endogenous Piezo1 with MNPs are related to the large size and structural complexity of the channel, as well as the limited availability of commercial antibodies (Ab) specifically targeting its mechanosensitive parts.

Here, we describe MagPiezo, a novel magnetogenetic toolkit for the remote and untethered manipulation of human endogenous Piezo1 channels in endothelial cells. We demonstrate for the first time that it is feasible to trigger the activation of Piezo1 channels without further cellular modification using a custom-made magnetic applicator and MNPs smaller than 20 nm engineered with anti Piezo1 antibodies for selective targeting. Through real-time calcium imaging, we show an increase in the cytosolic Ca^2+^ influx associated with magnetomechanical stimulation. Ca^2+^ entrance trigger different endothelial responses including cytoskeleton remodeling and transcriptional activation. This novel toolkit opens the venue for the direct manipulation of native Piezo1 ion channels in different cell types, presenting significant opportunities both for biological research and clinical applications.

## 2. Results

### 2.1 Design and characterization of MagPiezo, a novel magnetogenetic toolkit

The main aim of the work was to design and develop a magnetogenetic toolkit able to specifically target and remotely trigger the activation of endogenous human Piezo1 channels in endothelial cells. This toolkit, named MagPiezo, encompassed the design of two main components: i) small MNPs (< 20 nm) based on mixed ferrites (ZnMnFe2O4) with tunable magnetic anisotropy and functionalized with an antibody against the mechanosensitive domain of the Piezo1 channel; ii) a custom-made magnetic applicator designed to transmit torque to MNPs by controlling the evolution of magnetic field gradients along two perpendicular axes, thereby promoting the mechanical gating and opening of the channel (**Figure 1A**). One of the major challenges of using small MNPs is that they exert limited mechanical forces, in the fN to low pN range, which approaches the lower threshold of the channel gating (∼10 pN).[28] To overcome this limitation, we fine-tuned the chemical composition of the MNPs, specifically by introducing transition metals (M^2+^) in the spinel structure, to improve the magnetocrystalline anisotropy and thereby the resulting mechanical forces that they can exert under magnetic field stimulation. The MNPs used were mixed ferrites (Zn*x*Mn*y*Fe(3-*x-y*)O4, MF) synthesized via a one-step thermal decomposition method. In a previous study we investigated the effects of Mn^2+^ substitution (Mn*x*Fe*3-x*O4, *x*=0.14 up to 1.40) in the spinel structure using the same synthetic procedure, showing that Mn^2+^ incorporation allowed fine control of the anisotropy transition towards soft ferrite behavior.[29] To further improve the MFs effective magnetic moment, which is proportional to both saturation magnetization (Ms) and magnetic susceptibility,[24] we introduced an additional non-magnetic cation (Zn^2+^) into the spinel lattice. While non-magnetic cations preferentially occupy tetrahedral sites, strongly magnetic cations (Fe^2+^, Mn^2+^) are predominantly in octahedral sites, resulting in enhanced super-exchange interactions and improved Ms.[30–32] The resulting MFs were regular octahedral MNPs with average dimensions of 19 ± 2 nm x 26 ± 3 nm (**Figure 1B**). The one-step thermal decomposition method yielded highly reproducible nanoparticles coated with oleic acid (MF@OA), which could only be dispersed in organic solvents (**Figure S1**, Supporting Information). The transfer into the water was achieved using an amphiphilic polymer, poly- (maleic anhydride-*alt*-1-octadecene), PMAO, which was modified with a fluorescent molecule (TAMRA) to enable cellular tracking (MF@PMAO) (**Figure 1C**).[33–35] This polymer renders the MF water-soluble and leaves free carboxylic groups that can be used for further functionalization.

**Figure 1.**
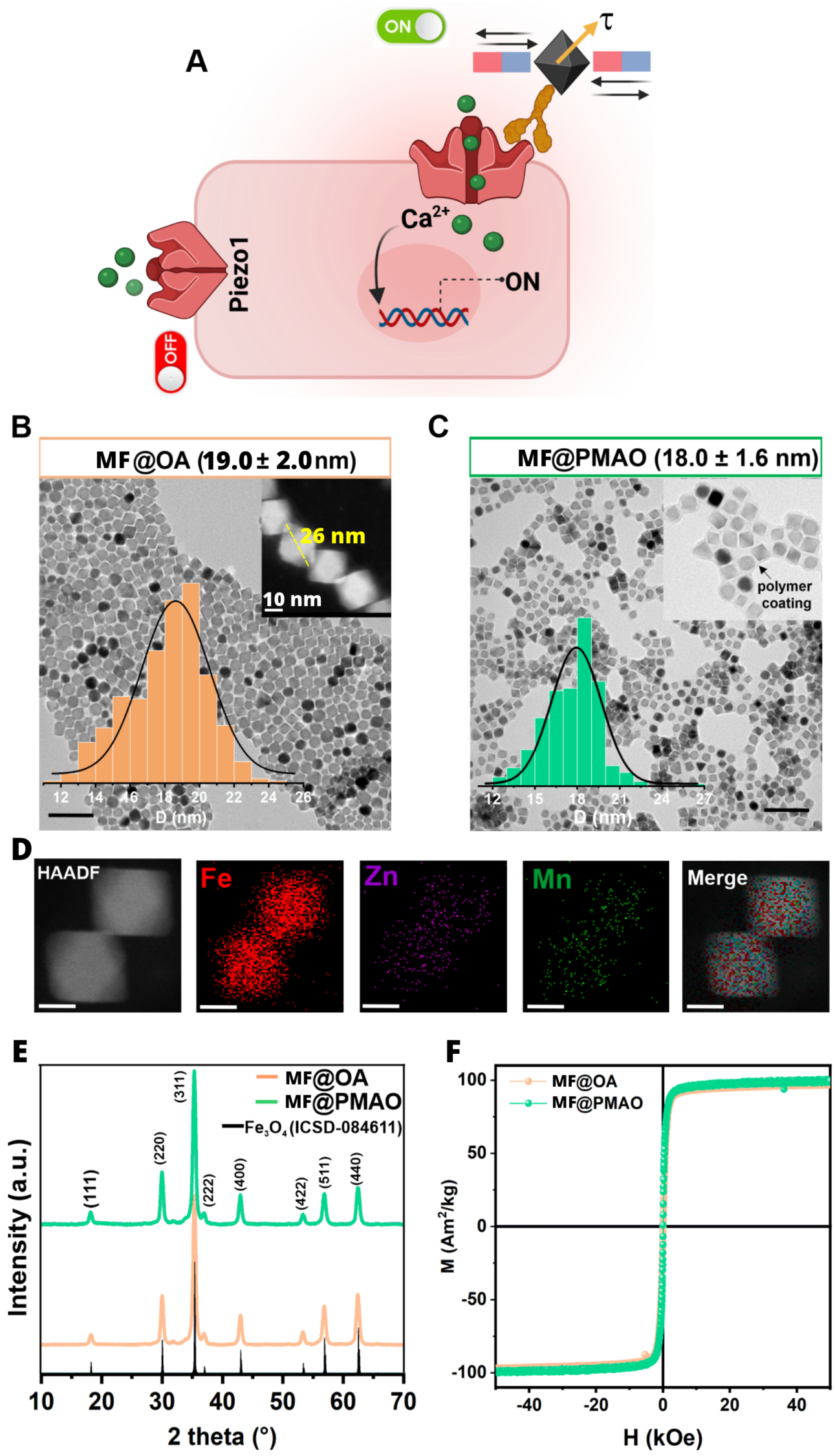
**A)** MagPiezo toolkit overview. **B**-**C)** TEM images of MF@OA and MF@PMAO MNPs, respectively (Scale bar =100 nm). Inset: Size distribution histograms fitted with a log-normal distribution. **D)** HAADF-STEM image and the corresponding EDX elemental mapping images of MF@OA MNPs (Scale bar =10 nm). **E**) Powder XRD patterns of MF@OA and MF@PMAO samples (the reference diffractogram corresponds to the Fe3O4 phase [ICSD 084611]). **F)** Hysteresis loops recorded at 290 K for MF@OA and MF@PMAO samples.

The elemental composition of the MF@OA system, measured by Inductively Coupled Plasma-Optical Emission Spectrometry (ICP-OES), was Zn0.29Mn0.18Fe2.53O4, and the presence of iron, zinc, and manganese across the octahedron was confirmed by High-Angle Annular Dark-Field Scanning Transmission Electron Microscopy (HAADF-STEM) coupled with Energy-Dispersive X-ray Spectroscopy (EDX) (**Figure 1D**). Both types of MNPs (MF@OA and MF@PMAO) exhibited single crystal structure indexing with the typical cubic spinel structure (ICSD-084611), with no detectable secondary crystalline phases, as can be seen from X-ray diffraction (XRD) patterns (**Figure 1E**). Both systems showed superparamagnetic (SPM) behavior at room temperature (RT) and a linear *M*(*H*) dependence on small fields. At RT, MF@PMAO system exhibited a maximum mass magnetic susceptibility value of 3.8 x 10^-6^ m^3^ kg^-1^, calculated as the slope of the *M*(*H*) curve in the low-field region (*H* < 100 Oe), indicating strong low-field magnetic responsiveness compared to typical MnFe2O4 or Fe3O4 nanoparticles.[36,37] The magnetization values were normalized to the organic content measured by thermogravimetric analysis (TGA, **Figure S2,** Supporting Information); a summary of the magnetic properties is provided in **Table S1**, Supporting Information. As expected, both MF@OA and MF@PMAO showed *Ms* values of approximately 100 Am^2^ kg^-1^, which are significantly higher than those we reported earlier for Mn*x*Fe*3-x*O4 of similar size (85 Am^2^ kg^-1^)[29] or other reported ferrites (typically <85 Am^2^ kg^-1^).[38–40] Importantly, the aqueous phase transfer using PMAO did not affect MF size, shape, crystalline structure or static magnetic properties, confirming the robustness of the coating approach (**Figure 1E-F**).

To enable remote magnetic stimulation, we designed a versatile magnetic applicator to be incorporated in a fluorescence microscope or to be used inside a cell incubator (**Figure 2A**). This setup allowed the application of homogeneous magnetic fields on a circular dish area (35 mm) where endothelial cells were seeded, enabling real-time calcium imaging or signaling pathways studies across hundreds of cells simultaneously. Initially, we attempted to implement a rotating Halbach magnet array, a configuration previously reported to open Piezo1 using large MNPs.[27] However, this configuration introduced mechanical vibrations in the cell culture, compromising the quality of real time calcium imaging data. We therefore developed a custom magneto-stimulation device, partially based on a previously reported system designed to deliver complex strain patterns to biological samples.[41] The device consisted of four independent sets of permanent magnets (10 x 10 x 3 mm thick, NdFeB) positioned symmetrically around a 35-mm circular sample area, with two sets of magnets aligned along each axis. Each magnet set was driven by independent servomotors, enabling precise control of both position and actuation frequency via an Arduino-based control system (**Figure 2B**). In our experiments, the device was configured to produce an alternating movement between two positions (denoted as A and B in **Figure 2C**) by simultaneously moving two opposing sets of magnets toward the center of the dish along one axis, while the other two sets moved outward along the perpendicular axis. (**Figure 2C** and **Video S1,** Supporting Information). This movement was then reversed in the next half-cycle, completing one full oscillation at a frequency of 1 Hz.

**Figure 2.**
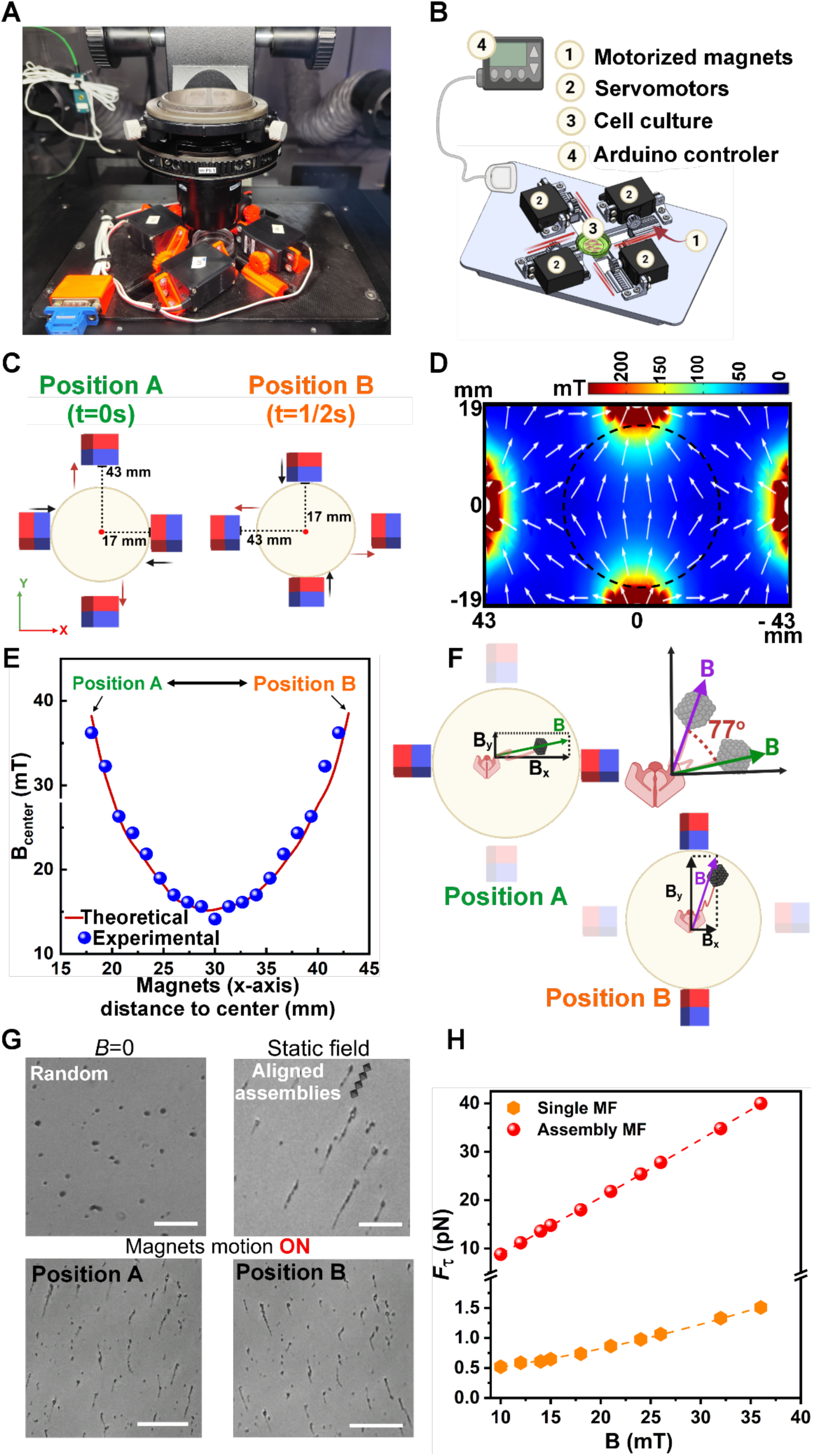
Design and characterization of a magnetic applicator for torque generation. **A)** Custom-made magnetic applicator setup incorporated in a fluorescence microscope. **B)** Components of the magnetic field applicator. **C)** Dynamic magnetic field is achieved by simultaneously switching the sets of magnets from Position A to B at a frequency of 1 Hz. Magnets move along their axis from the closest position to the dish (17 mm from the center) to the farthest position (43 mm from the center). **D)** Surface magnetic flux density map generated by the magnetic applicator at the fixed position B (white arrows represent normalized field magnitude). **E)** Theoretical (red line) and experimental (blue dots) validation of the magnetic field intensity measured at the center of the petri dish during the full motion of the magnets from position A to position B. **F)** Schematic representation of the rotational movement generated by the magnetic applicator. **G)** Dynamics of MF MNPs assembly formation and induced rotational motion: MF MNPs under zero magnetic field, under experiencing initial static magnetic field, and upon activation of the alternating motion of the permanent magnets. Scale bar: 50 μm. **H)** Comparison of torque forces generated by a single superparamagnetic MF MNP versus an assembly (10^4^ particles) depending on the intensity of the applied field. Polynomial fittings (dashed lines) are applied to both datasets to highlight the quadratic dependence of torque force with the applied field intensity.

The intensity of the magnetic field was modulated by varying the number of magnets included in each set. To do so, the surface magnetic flux density was theoretically simulated using COMSOL Multiphysics, considering sets of 2, 4, 6, 8 or 10 permanent magnets (**Figure S3** in the Supporting Information). While the addition of magnets to the configuration resulted in an increase in the strength of the magnetic field, this enhancement became less pronounced beyond the inclusion of six magnets. Additionally, from a practical point of view, the onset of mechanical vibration was observed at this point. Consequently, six permanent magnets per set were selected, as they generated the highest magnetic field without introducing undesirable mechanical vibrations.

Subsequently, we simulated the surface magnetic flux density generated by the whole magnetic applicator (4 sets of 6 permanent magnets each) at a fixed position (position B) (**Figure 2D)**. This graph density shows that the field lines are oriented along the axis where the magnets are closest. The magnetic strength along the X-axis ranges from 38 mT at the centre to 250 mT near the edges, forming a quadrupolar distribution with a clear outward gradient (**Figure 2D)**. To later include four simultaneous experimental conditions, a cell culture insert of approximately 17 mm diameter was placed in the center of the dish. Within this region, the magnetic field was relatively uniform, with intensities ranging from 38 to 60 mT and low gradients (<15 mT mm^-1^), allowing it to be considered as a homogeneous field (**Figure S4** in the Supporting Information). Lastly, the alternating motion of the magnetic applicator between positions A and B generated a dynamic magnetic field at the center of the dish, ranging from 15 to 38 mT. These values were validated through experimental measurements using a Gaussmeter placed at the center of the dish (see experimental section), yielding values of 14-36 mT, in strong agreement with simulation data (**Figure 2E**). Moreover, the position and motion of the magnets were designed to induce a rotational movement of the magnetic field of approximately 77° (**Figure 2F**).

To calculate the forces generated by the MF MNPs, it is important to note that in the absence of an external magnetic field, SPM nanoparticles undergo random Brownian motion, and thus the direction of the magnetic moments fluctuates rapidly, resulting in no net magnetization. When a static magnetic field is applied, the magnetic moments of the SPM nanoparticles tend to align with the field direction, promoting the formation of linear assemblies or chain-like aggregates along their magnetic easy axes due to dipolar interactions. In our case, this prealignment occurs when the MF MNPs are placed in the magnetic field applicator, prior to initiating the alternating motion of the set of magnets (**Figure 2G** and **Video S2** in the Supporting Information). This self-assembling of MNPs into chains generated in situ by a static magnetic field has been previously used to enhance the mechanical torque produced in cells when using a rotating magnetic field.[42] Thereafter, upon activation of the alternating motion of the permanent magnets, the direction of the magnetic field vector changes dynamically. Under these conditions, the MNP assemblies previously formed attempt to realign their magnetic moments with the rotating field, generating a magnetic torque (**Figure 2G)**.[43] Importantly, this phenomenon can be observed in both aqueous and viscous environments (**Video S3** in the Supporting Information).

With all this in mind, we first estimated the mechanical force acting on a single SPM nanoparticle under a maximum magnetic field intensity of 36 mT (see **Section 3.1** in the Supporting Information). In the absence of interparticle interactions, each nanoparticle experiences a magnetic torque of *τ*_1_ = 2.0 ×10^-20^ Nm, which translates into a torque force of *Fτ* =1.5 pN. As shown in **Figure 2D**, the mechanical force due to magnetic torque *F*_τ_ shows a dependence on the square of the magnetic field intensity (*B*²), consistent with the torque scaling as *τ* ∝ *B*² at low field regimes. Moreover, by using the Langevin function, we calculated the dimensionless parameter *χ* (which compares magnetic energy to thermal energy). The computed *χ* = 14 indicates that the magnetic moment is strongly, though not perfectly, aligned with the applied magnetic field.[44] Thus, we calculated an average angle *θ* of approximately 21° between the magnetic moment and the magnetic field. This misalignment is critical for torque generation by our SPM nanoparticles, suggesting that even individual MF MNPs could generate sufficient force to gate Piezo1 channels.

Next, we estimated the torque experienced by a single assembly of aligned MF MNPs with an average length of 3 μm and width 0.2 μm (**Figure 2H** and **Section 3.2** in the Supporting Information). The torque per assembly was *τ_assembly_* = 2.6×10^−16^ Nm, indicating that one MF assembly can generate 9 - 40 pN at 10 - 36 mT. As shown in **Figure 2H**, the torque force in MF assemblies scales with the applied field more rapidly than for individual MF MNPs, highlighting the enhanced mechanical output of collective assemblies.[43]

### 2.2 Oriented conjugation of anti-Piezo1 Ab on MF surface for selective targeting of endogenous Piezo1 in endothelial cells

As already mentioned, to date the few reports dealing with the specific gating of Piezo1 channels with magnetic particles use non-endogenous receptors. In all cases the channel was exogenously expressed and modified with specific tags that were used to attach the MNPs.[25–27] To open the endogenous channel using small MNPs, it is crucial to couple the MNPs with an anti-Piezo1 antibody (AbPIEZ) able to recognize not only the extracellular portion of Piezo1, but also part of the mechanical region (2199-2426 aa in Cap region). [28] Furthermore, the Ab orientation on the MNP surface is a critical yet sometimes overlooked aspect. Given the complex three-dimensional structure of Abs, it is essential that, upon immobilization, their antigen-binding (Fab) regions remain available for epitope recognition. While random functionalization strategies often result in a reduction in the antigen recognition efficiency,[45] the orientation of the Ab based on the kinetic differences between ionic adsorption and covalent attachment, ensures a correct positioning (**Figure 3A**).[46,47] At the same time, it avoids the need for adaptor molecules or Ab modification, which can be costly or lead to partial loss of Ab activity.

**Figure 3.**
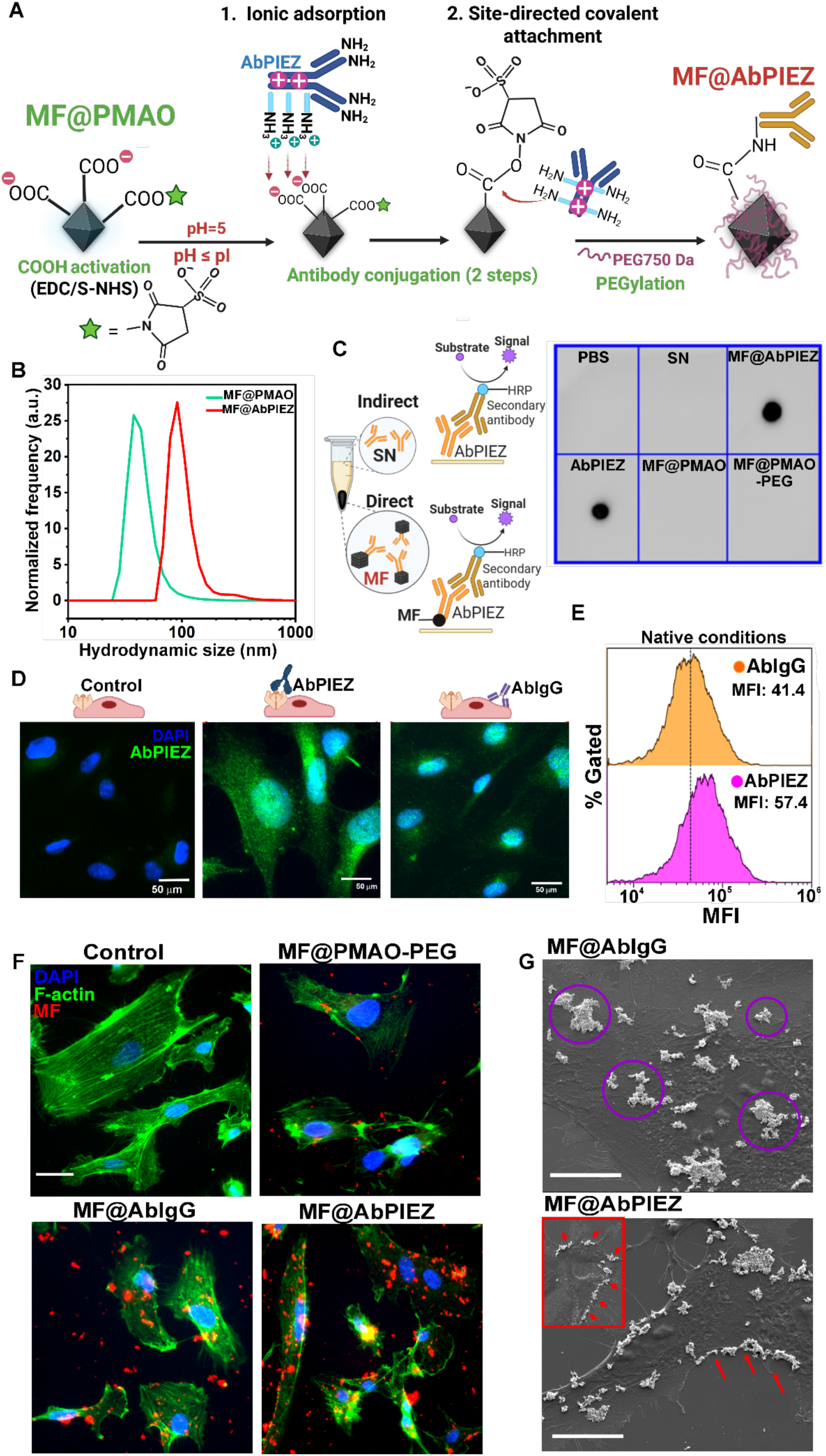
Engineering MF MNPs for selective targeting of Piezo1 channel. **A)** Schematic representation of AbPIEZ oriented conjugation onto the MF surface following a two-step methodology and their subsequent PEGylation (MF@AbPIEZ). **B)** Hydrodynamic sizes (in number) of MF@PMAO and MF@AbPIEZ. **C)** Dot blot immunoassay showing the successful functionalization of the MFs with AbPIEZ. The schematic representation depicts two ways of indirectly assessing that the Ab was attached to the MF: i) MF@PMAO, MF@PMAO-PEG or MF@AbPIEZ were deposited on the membrane or ii) the SN of the functionalization process was analyzed and compared to the Ab that was added (100%). **D)** Comparisons of live immunofluorescence images of HUVEC cells expressing native Piezo1 and treated with AbPIEZ or isotype control Ab (AbIgG) (n = 2 biological replicas). Scale bar: 50 μm. **E)** Flow cytometry histograms of HUVEC cells incubated with AbPIEZ or its isotype control (AbIgG), followed by a rabbit anti-IgG secondary antibody. Geometric Mean Fluorescence Intensities (MFI) are reported for each sample (x10³). **F)** Fluorescence microscopy images of HUVEC cells without MFs (Control) or treated for 40 min with unconjugated MF (MF@PMAO-PEG), MF@AbIgG, or MF@AbPIEZ. F-actin is stained in green, nuclei in blue and MF in red (TAMRA) (n =2 biological replicas). Scale bar: 20 μm. **G)** SEM images of HUVEC cells treated with MF@AbIgG or MF@AbPIEZ nanoparticles highlighting their distinctive spatial distribution on the cell surface. Scale bar: 20 μm.

Based on our previous works,[47,48] we proposed a functionalization protocol that involves an initial rapid ionic adsorption of the Abs onto the negatively charged MF surface, conducted at a pH where the Abs are positively charged. For polyclonal AbPIEZ, with an isoelectric point (pI) between 5.2 and 8.5, ionic pre-adsorption was performed at a pH of 5 (**Figure S5** in the Supporting Information). Since the Ab is adsorbed through its richest positive zone (typically the constant region, Fc),[47] the Fab region remains exposed, and the antigen binding capacity is preserved. Once the Ab is oriented, the second step implies a slower covalent attachment of the Ab to the MF, that will occur only where the first ionic adsorption took place. For this step, the carboxylic groups on the MF surface were previously activated using carbodiimide chemistry. Once the Ab was covalently attached, the remaining activated carboxylic groups were passivated using PEG molecules (MF@AbPIEZ).

Throughout the surface modification process, the hydrodynamic sizes of MF@PMAO increased, indicating successful AbPIEZ conjugation without causing aggregation (**Figure 3B**). The success of the coupling reaction was verified by a dot blot immunoassay, analyzing both the supernatant (SN) from the functionalization process and the MF@AbPIEZ (**Figure 3C**).[35,49] In all cases, the intensity of the signal dot is directly proportional to the amount of Ab present in the sample. The SN from the functionalization process was compared to control samples containing the same amount of added Ab, representing 100% of the input AbPIEZ. Complete Ab binding to the MFs was confirmed, as no signal was detected in the SN. Additionally, we also analyzed the MF with Ab (MF@AbPIEZ) and we used as controls MF@PMAO and MF functionalized with PEG but lacking the Ab (MF@PMAO-PEG). Once deposited on the membrane, MF@AbPIEZ produced a strong signal caused by the Ab attached to the MF, whereas MF@PMAO or MF@PMAO-PEG did not give any signal. Based on the size and morphology of the MF nanoparticles and the amount and size of AbPIEZ conjugated, we estimated an average of two AbPIEZ molecules per nanoparticle (**Section 4** in the Supporting Information). Higher antibody loading was avoided, as it compromised MF colloidal stability.

Before assessing if MF@AbPIEZ could recognize endogenous Piezo1 in cells, we verified the presence of this channel in HUVEC cells using the same Ab employed to functionalize the MFs. To ensure that we were exclusively detecting the extracellular domain of Piezo1, we performed a live cell immunofluorescence assay, where the Ab recognizes the epitope without the prior use of fixatives or permeabilization agents. Under these native conditions, a strong labelling of Piezo1 was observed, while a faint basal signal was detected using an isotype Ab as a negative control (**Figure 3D**). This result was corroborated by flow cytometry, which also allowed comparison between surface expression of Piezo1 (native conditions) and total cellular level (permeabilization conditions) (**Figure 3E** and **Figure S6** in the Supporting Information). We next evaluated whether MF@AbPIEZ could selectively recognize endothelial HUVEC cells, incubating the cells with MF@AbPIEZ at 50 μg Fe mL^-1^ for 40 minutes. As negative controls, we included MF@PMAO-PEG and MF functionalized with the isotype IgG antibody (MF@AbIgG). Fluorescence microscopy revealed a stronger signal in cells treated with MF@AbPIEZ compared to those treated with controls (**Figure 3F**). In the case of MF@AbIgG we also observed a fluorescence signal, which could be related to an unspecific recognition mediated by the IgG, confirming the immunolabelling and flow cytometry results (**Figure 3D**). Noteworthy, MF@PMAO-PEG treated cells showed no detectable signal, confirming that the conjugation with Abs is necessary for Piezo1 targeting. To further explore the interaction of MF@AbPIEZ with HUVEC, cells were incubated with the particles for 40 minutes, fixed, and analyzed by Scanning Electron Microscopy (SEM). Consistent with previous observations, SEM images revealed that most of the MF@AbPIEZ and MF@AbIgG systems were not internalized by HUVEC cells after 40 minutes of incubation, likely limiting their action to the cell membrane (**Figure 3G**). However, we found a very distinctive spatial distribution of the MFs depending on the Ab used in the functionalization process. While MF@AbPIEZ appeared predominately along the edges of the cells, the MF@AbIgG were distributed all over the cell in a more random fashion, suggesting an unspecific interaction. These results indicate that while both MF systems can interact with endothelial cells, specific and directional binding is achieved only through a successfully selected antibody-mediated targeting.

### 2.3 Real-time calcium dynamics after magnetomechanical stimulation

Prior to testing the remote magnetomechanical activation of Piezo1, we assessed the cytotoxicity of the MF@AbPIEZ alone and in combination with magnetic field. Cells were treated with the MF MNPs, exposed to a sustained 40-minute magnetic field, and stained with calcein-AM and ethidium homodimer-1. No cell death was observed, as shown in **Figure S7A**, Supporting Information. The TUNEL assay performed 24 hours after exposure to MF alone or in combination with magnetic stimulation also confirmed the lack of apoptotic cells (**Figure S7B,** Supporting Information).

Interestingly, when we magneto-stimulated the cells for 30 minutes, aligned assemblies of FM NPs (as observed in water and glycerol) were found at different locations on the cell membrane, which conserved their rotational motion locally (**Video S2** in the Supporting information). As the total torque applied to the whole cell increases proportionally with the number of assemblies grafted,[43] the presence of multiple actively rotating assemblies should amplify the global mechanical torque exerted on the entire cell.

We then determined calcium dynamics over time to evaluate in real-time the possibility of Piezo1 gating by our magnetogenetic toolkit (**Figure 4A-i**). For magnetomechanical stimulation, cells were treated with MF MNPs for 40 minutes and incubated with Fluo-4 AM before being stimulated with the magnetic applicator for 30 minutes. Fluorescence imaging of Ca^2+^ dynamics was acquired during the whole duration of the magnetomechanical stimulation, allowing for continuous tracking of individual cells. Image acquisition started 1.5 minutes before applying the alternating motion of permanent magnets, and the average fluorescence intensity of each cell during this period was considered as their baseline fluorescence (F0) (**Figure 4A**-**ii**). As already reported for neurons stimulated with nanomagnetic forces,[50] a continuous Ca^2+^ influx could be observed over time during the magneto-stimulation of cells treated with FM@AbPIEZ (**Figure 4A**-**iii)**. These ΔF/F0 Ca^2+^ traces can be decomposed into a resting Ca^2+^ waveform, enabling both the quantification of the cumulative Ca²⁺ flux during stimulation (calculated as the area under the resting waveform) and the detection of Ca^2+^ transients or events (**Figure 4A**-**iii**). [50] To do that, we developed MagCalcium,[51] a dedicated code to track both calcium events per cell and continuous Ca^2+^ influx (see **Section 6**, Supporting information).

**Figure 4.**
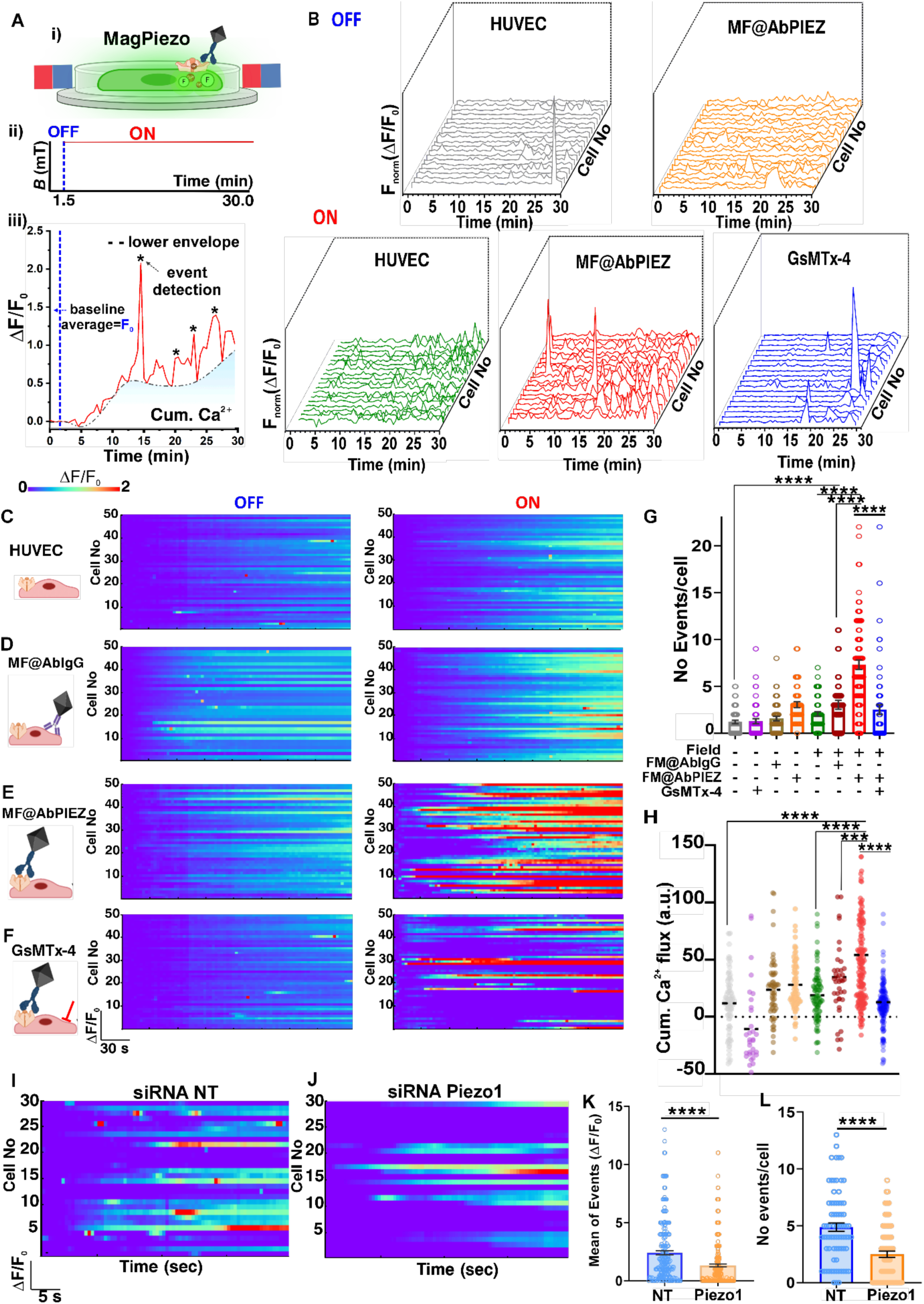
Calcium dynamics in HUVEC cells. **A)** i) Schematic representation depicting real-time calcium imaging; ii) Experimental conditions used for sustained magnetomechanical stimulation. iii) Our Ca^2+^ flux analysis combines estimation of the continuous Ca^2+^ flux (cumulative Ca^2+^) during magnetomechanical stimulation period and events detection. Cumulative Ca^2+^ was measured as the finite sum of the area under the resting Ca^2+^ waveform during the stimulation period. To obtain the resting Ca^2+^ waveform we fit a lower envelope to the signals (ΔF/F0) with the envelope function implemented in the MagCalcium code. The somatic events were considered as peaks with a prominence of both > 5% ΔF/F0 and 3 standard deviations above F0. **B)** Representative normalized calcium fluorescence (Fnorm) intensity traces (ΔF/F0) for different conditions (20 cells each condition), stimulated (ON) or not (OFF) with magnetic field. To normalize the signal, the lower envelope was subtracted from the ΔF/F0 traces. **C-F)** Heatmaps of calcium dynamics (representing ΔF/F0) of cells subjected to different treatments (50 cells each condition), not stimulated with magnetic field (OFF) or under sustained magnetic stimulation (ON): C) Control cells (without MNPs), D) MF@AbIgG -treated cells, E) MF@AbPIEZ -treated cells and F) cells treated with MF@AbPIEZ and previously blocked with the GsMTx-4 inhibitor. **G)** Quantification of number of events per cell (n = 4 biological replica). Statistical analysis was performed using one-way ANOVA and Holm-Sidak’s multiple comparisons (mean of each treatment against the mean of MF@AbPIEZ + field). **H)** Cumulative Ca^2+^ flux for the different conditions tested (n = 4 biological replica). Statistical analysis was performed using one-way ANOVA and Dunet multiple comparisons test (mean of each treatment (dashed lines) against the mean of MF@AbPIEZ + field). **I-J)** Heatmaps of calcium dynamics representing ΔF/F0 of cells previously treated with (**I**) NT siRNA (siRNA control) or with **(J)** siRNA Piezo1 (n = 2 biological replica). After siRNA treatment, both samples were incubated with MF@AbPIEZ and exposed to the magnetic field. **K-L)** Quantification of average of Ca^2+^ events (n = 219 and 242 cells) and number of events per cell (n=74 and 82 cells) of the cells treated with NT siRNA and siRNA Piezo1, respectively. Statistical analyses were performed using a two-tailed Student’s t test. Data are shown as mean ± SEM.

Initial observations suggest that MagPiezo induced more frequent Ca²⁺ transients in HUVEC cells than either MF@AbPIEZ or magnetic field applications alone (**Figure 4B** and **Video S4** in the Supporting Information). This effect was almost completely abolished by pretreatment with the Piezo1 inhibitor GsMTx4, confirming the specificity of the response (**Figure 4B** and **Video S4** in the Supporting Information). Notably, the calcium traces induced by MagPiezo could be observed during the whole duration of the magneto-mechanical stimulation, differing from those produced using the agonist Yoda1 (5 μM), which triggered a global and rapid Ca^2+^ influx response (**Figure S8**, in the Supporting Information).

To assess whether MagPiezo can modulate calcium responses depending on the duration of magnetomechanical stimulation, we compared cells stimulated for 10 and 30 minutes. We found that longer stimulation times led to a proportional increase in both the number of calcium events and the overall calcium accumulation (**Figure S8** in the Supporting Information). These findings highlight that this mode of mechanical stimulation does not alter the homeostatic calcium dynamics of HUVEC cells. In astrocytes, for instance, the use of sustained mechanical stimulation implemented with elastomers and magnets led to a pronounced loss of calcium oscillations after a few minutes of stimulation.[41]

Thereafter, we generated heatmaps reflecting the total cellular fluorescence traces (**Figure 4C-F**) and quantified both the number of events per cell (**Figure 4G**) and Ca^2+^ accumulation during the magneto-stimulation period (**Figure 4H**). In agreement with **Figure 4B**, neither MF@AbPIEZ nor magnetic field alone generated significant changes in the number of calcium events or in the cumulative Ca^2+^ (**Figure 4C, 4E,4 G and 4H**). To further provide validation of the selectivity of MagPiezo, we treated cells with MF@AbIgG under the same experimental conditions, showing no significant Ca^2+^ influx over time (**Figure 4D**, **4G**, **4H** and **Video S5** in the Supporting Information), highlighting the importance of selective membrane targeting for magnetomechanical modulation. Cells previously treated with the GsTMx-4 inhibitor resulted in a significant decrease in the number of events and calcium accumulation (**Figure 4B, 4F, 4G and 4H**).

To fully demonstrate the role of Piezo1 in these responses, we used siRNA to knockdown Piezo1 expression in HUVEC cells. The efficiency of the siRNA treatment was evaluated by quantitative real-time PCR (RT-qPCR) and showed a significant reduction in Piezo1 mRNA levels. Specifically, Piezo1 expression was reduced by 46% and 88% at 48- and 72-hours post-treatment. As it can be seen in **Figure 4I-L**, both the mean intensity in calcium influx and the number of events per cell significantly decreased (by 54% and 51%, respectively) in the cells where Piezo1 was knocked down compared with cells treated with a non-targeting (NT) siRNA (**Video S6** in the Supporting Information), highlighting the role of this MS ion channel in mediating Ca^2+^ responses in endothelial cells.

### 2.5 MagPiezo-mediated stimulation induces cellular responses at different levels

Next, we examined whether the intracellular calcium influx, induced by Piezo1 gating following magnetomechanical stimulation, modulated different downstream biological responses. For this we focused on the established responses from endothelial cells as they adapt to mechanical forces, such as shear stress.[52].

#### 2.5.1 Magnetomechanical activation of Piezo1 drives calcium-dependent cytoskeletal remodeling

The organization and dynamics of the actin cytoskeleton are crucial for maintaining cell shape and tissue integrity for enabling cellular adaptation to mechanical input. In addition, the actin cytoskeleton is responsive to changes in cytosolic calcium concentration[53] and is known to interact with Piezo1.[54–56] To determine the effect of MagPiezo on actin remodeling, we labelled cells with fluorescently conjugated phalloidin, a probe that labels filamentous actin (F-actin). In unperturbed control cells, the actin cytoskeleton is organized mainly as a continuous cortical meshwork and thicker cables or stress fibers that span and mechanically support the cell (**Figure 5A**, see inset). The addition of MF@AbPIEZ strengthened some of the actin cables, displayed as a slight increase in fraction of highly intense actin cables (**Figure 5A** and **B**, see experimental section). Interestingly, this effect did not change with 20 minutes of sustained magnetomechanical stimulation, while the fraction of highly intense actin cables was completely abolished when Piezo1 calcium influx was blocked by using GsTMx-4 (**Figure 5A** and 5**B**). We then quantified the redistribution in F-actin by calculating the ratio between actin accumulating near the cell edges versus the actin present at the cell interior (**Figure 5C**). The effect of magnetomechanical stimulation was prominent in enhancing the F-actin cables at the cell edges and cell-cell contacts (**Figure 5A** and 5**C**). The stimulation resulted in an almost 40% increase of the F-actin accumulation at the cell edges, likely involving the strengthening of cell-cell contacts (**Figure 5D**). Actin remodeling induced by the magnetomechanical stimulation of Piezo1 presumably involves myosin-mediated contractility. In fact, one of the immediate consequences of increased myosin activity is the reinforcement of stress fibers.[57] Consistent with this, we observed that stimulated cells exhibited the largest reduction in spread area from 920 ±395 µm^2^ to 549±216 µm^2^.

**Figure 5.**
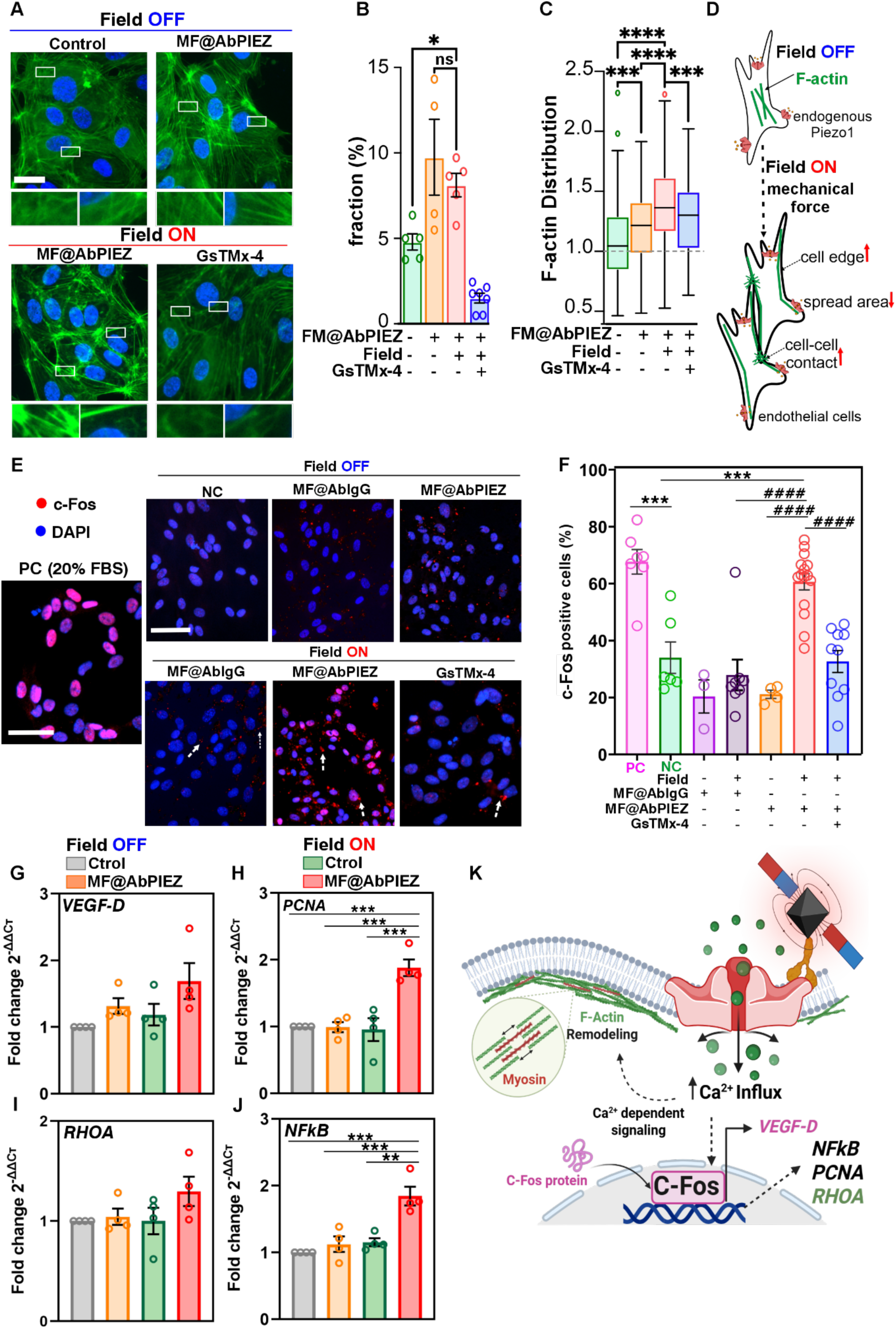
Biological responses resulting from MagPiezo application in endothelial cells. **A)** Fluorescence images of cells stained for the nucleus (blue) and F-actin (green). Each panel displays a representative image of cells that have undergone the indicated treatment (n = 2). Zoomed-in panels display the fibers situated around the cell interior and the edges. Scale bar 50 μm. **B)** Fraction of strengthened actin fibers in cells from different treatments. **C)** Distribution of filamentous actin within the cells, where <1 indicates an accumulation within the cell interior and >1 indicates a strengthening at the cell edges. **D)** Schematic representation of the proposed F-actin redistribution mechanism following magnetomechanical stimulation. **E)** Representative immunofluorescence images showing c-Fos translocation to the nucleus in endothelial cells following magnetomechanical stimulation (n = 3). c-Fos (red) and nuclei stained with DAPI (blue). MF MNPs are indicated with white arrows. Scale bar 100 μm. **F)** Percentage of c-Fos^+^ cells across treatments, normalized to the total number of analyzed cells (n=1063 (PC), 880 (NC), 344 (MF@AbIgG), 483 (MF@AbPIEZ), 875 (MF@AbIgG+field), 3072 (MF@AbPIEZ + field), 2844 (GsTMx-4). Cells were classified as positive when the Pearson correlation coefficient (P) exceeded 0.35 using Cell Profiler program. Data are shown as the means of individual technical replicates ± SD. Statistical significance was determined by one-way ANOVA followed by Tukey test. (#) symbol is used for comparisons performed against the MF@AbPIEZ + field sample and (*) for comparisons performed against the positive control (PC). **G-J)** Relative mRNA expression levels of key genes involved in calcium signaling pathways obtained 20 minutes after magnetic stimulation (n = 4): (F) *VEGF-D*, (G) *RHOA*, (H) *PCNA*, and (I) *NFκB*. **K)** Graphical summary illustrating the key endothelial responses induced by MagPiezo. Data are shown as mean ± SEM

#### 2.5.2 c-Fos Activation as a readout of MagPiezo-mediated mechanotransduction

c-Fos is a transcription factor that plays a central role in mediating nuclear responses to signals acting at the cell membrane. For instance, shear stress in endothelial cells leads to a rapid induction of the proto-oncogen c-Fos, that upon binding with c-jun forms a transcriptional activator of various genes.[58] Similarly, nuclear c-Fos levels can be elevated in response to mechanical stimuli such as magnetomechanical forces and ultrasounds.[27,59] In our study, nuclear c-Fos accumulation was analyzed immediately following magnetic field stimulation (**Figure 5E**). As a positive control (PC), we added FBS to cells previously serum-starved for 24 h. In this case, up to 66% of the cells accumulated c-Fos in their nuclei (**Figure 5E** and **5F**). Importantly, cells that were stimulated using MF@AbPIEZ in combination with a sustained magnetic field exposure (field ON) had a nuclear accumulation of c-Fos in 61% of the cells, a level comparable to the PC. The fact that nuclear c-Fos level was crucially dependent on the application of the magnetic field highlights the robust efficacy of the MagPiezo system in activating mechanosensitive transcriptional responses. In stark contrast, the nuclear accumulation of c-Fos in cells treated with MF@AbIgG was comparable to that of the negative control condition and marginally increased in response to the application of a magnetic field (**Figure 5E** and **F**). Furthermore, pretreatment with the Piezo1 inhibitor GsTMx-4 prevented nuclear c-Fos accumulation. Altogether these data demonstrate that the observed transcriptional response of c-Fos induced by our toolkit, as measured though its nuclear accumulation, is specific and Piezo1-dependent.

#### 2.5.3 Downstream effects of Piezo1 activation through MagPiezo

As a third functional consequence of Piezo1 gating using our MagPiezo toolkit, we focused on changes in gene expression of four key components in the physiological response of endothelial cells. Using RT-qPCR we followed the transcriptional levels of nuclear factor kappa-light-chain-enhancer of activated B cells (*NFkB*), proliferating cell nuclear antigen (*PCNA*), Ras homolog family member A (*RHOA*) and vascular endothelial growth factor D (*VEGF-D*) (**Figure 5G**-**J**). These genes are implicated in several cellular processes, including inflammation, proliferation and apoptosis, and can be modulated in response to Piezo1 activation.[60,61]

Magnetomechanical stimulation of Piezo1 was performed as explained before, and after 20 minutes RNA was extracted, and transcript levels were determined. The transcripts for *VEGF-D* and *NFkB* were upregulated marginally by the presence of MF@AbPIEZ only (**Figure 5G** and **J)**. Importantly, and consistent with the above presented data, the expression of all 4 genes increased significantly and exclusively when the cells were treated with MF@AbPIEZ and stimulated using the magnetic field (**Figure 5 G**-**J**). The increase in *RHOA* expression agrees with the hypothesis that the functional changes in actin organization are due to increased myosin activity. The increased expression levels of *PCNA*, *NFkB* and *VEGF-D* on the other hand suggest enhanced activation of angiogenesis and endothelial cell proliferation processes.

Altogether these results demonstrate that Piezo1 opening using MagPiezo toolkit directs c-Fos to the nucleus, rearranges the actin cytoskeleton and directly affects the transcript levels of key genes in endothelial cells (**Figure 5K**).

## 3. Discussion

In this study, we introduce MagPiezo, a novel magnetogenetic platform that enables the remote activation of native Piezo1 channels in human endothelial cells without the need for genetic modification. Our results demonstrate that sustained magnetomechanical stimulation using engineered MF MNPs elicits precise endothelial responses, underscoring the potential of this approach to interrogate and control Piezo1 mechanotransduction under both physiological and pathological vascular conditions.

A key innovation of this work lies in the direct use of 19 nm MF nanoparticles to generate wireless torque forces in the piconewton range sufficient to activate Piezo1 when using a customized magnetic applicator that can alternate the motion of two sets of magnets. This combination represents a novel methodology to trigger human endogenous Piezo1 channels opening; noteworthy, the force generated can activate Piezo1 but does not reach the threshold reported for causing membrane damage (hundreds of pN).[62] Compared with previous studies where the mechanical opening of Piezo1 was smartly demonstrated in neurons both in vitro and in vivo using larger particles, [25–27] MagPiezo stands as a simpler alternative, as MF MNPs do not need to be further incorporated onto larger beads of 200 or 500 nm. This avoids additional chemical steps and reduces the risk of non-specific targeting.[62] Moreover, the direct targeting of the MFs to the mechanosensitive domain of Piezo1 also contrasts to previous strategies that relied on genetically modified ion channels.[25,26,28] MagPiezo acts directly on endogenous Piezo1, preserving native cellular physiology and mimicking physiological conditions.

Finally, the possibility of gating mechanoreceptors using low field intensities (< 40 mT) and frequencies, paves the way for an easy translation into an in vivo system. This setup is highly customizable, allows integration with fluorescence microscopes, circumvents the need for bulky and costly power sources, relying instead on low-intensity magnetic fields that are easy to generate, does not require expensive or large-scale equipment, and does not interfere with other biological processes. Moreover, the use of low-frequency actuation minimizes inelastic energy dissipation, effectively preventing undesirable heat generation.

Mechanistically, MagPiezo activation of Piezo1 induces a cascade of intracellular events, including calcium influx, cytoskeletal reorganization, and transcriptional activation. Our findings demonstrate that magnetomechanical stimulation using MagPiezo selectively activates Piezo1 channels, promoting an increase in both the number of calcium events per cell and the calcium accumulation flux, without disrupting calcium homeostasis dynamics. In contrast, cells treated with MF@AbIgG exhibited only a negligible increase in calcium influx, despite the unspecific attachment of these MNPs to the cell membrane. This underscores the critical role of Piezo1 in targeting effective modulation. Furthermore, the significant reduction in the calcium influx and events observed in cells where Piezo1 was silenced by siRNA, clearly confirms the specificity of our system and supports the involvement of this MS ion channel in the responses.

One of the most striking effects observed was the reorganization of the actin cytoskeleton. Within only 20 minutes of stimulation, we detected a marked increase in F-actin cable intensity, particularly at the cell cortex, suggesting enhanced cytoskeletal tension and remodeling. This effect was abolished by inhibition of Piezo1-mediated calcium entry, reinforcing the mechanistic link between calcium signaling and actin dynamics.[63,64] These findings align with previous studies demonstrating the role of Piezo1 in cytoskeletal regulation via calcium-dependent signaling pathways.[56] Additionally, the reduced cell spread area observed upon stimulation is consistent with increased myosin contractility and stress fiber reinforcement, further supporting activation of mechanotransduction pathways.

Concomitantly, we observed robust nuclear accumulation of the transcription factor c-Fos only in cells treated with MagPiezo. This response mirrors other transcriptional programs activated by mechanical stimuli, including magnetomechanical and ultrasonic forces.[27,59] Thereafter, c-Fos can directly induce the expression of *VEGF-D*, which is related with angiogenesis.[65] On the other side, other downstream signaling pathways involved in proliferation (e.g., via *PCNA* or *NFkB*) and cytoskeletal remodeling (e.g., via *RHOA*) could be activated by MagPiezo. For instance, while c-Fos does not directly regulate *NFkB*, both transcription factors can be co-activated by shared upstream signals such as calcium influx and MAPK activation in response to mechanical stress, inflammation, or osmotic changes.[66] These findings suggest that MagPiezo not only triggers immediate responses like calcium entrance, but also induces gene changes with important implications for endothelial biology.

Compared to conventional mechanical stimulation techniques, such as fluid shear stress, optical tweezers, or AFM, MagPiezo offers several advantages. It is non-invasive and it can activate multiple cells at the same time, enabling studies of cellular mechanics with high spatial resolution. Interestingly, other mechanosensors such as Notch, integrins or cadherins fall within a similar opening range that Piezo1, probing the versatility of MagPiezo. [14] Moreover, the integration of the magnetic field applicator in the microscope allows for real-time monitoring of calcium dynamics.

In conclusion, MagPiezo represents a significant advancement in the field of magnetogenetics, providing a powerful and specific toolkit for remotely controlling native MS ion channels without genetic manipulation and using low intensity-frequency magnetic field. This platform lays the groundwork for future therapeutic strategies in regenerative and cardiovascular medicine, where precise modulation of endothelial function could enhance tissue regeneration, angiogenesis, and the regulation of vascular inflammation. Beyond vascular biology, MagPiezo’s modular design enables adaptation to other mechanically responsive cell types, including neurons, immune cells or stem cells.

## 4. Experimental Section

### Synthesis of mixed ferrite ZnMnFe2O4 (MF@OA)

through one-step thermal decomposition method: In brief, 3.33 mmol of Fe(acac)3, 1 mmol of Mn(acac)2, and 0.67 mmol of Zn(acac)2 were placed in a three-neck flask with of oleic acid (15 mmol, 5.29 mL) as a surfactant, 10 mmol, 2.58 g of 1,2-hexadecanediol in benzyl ether (50 mL). The mixture was mixed and mechanically stirred (100 rpm) under vacuum for 30 min at room temperature (RT) and then, under a flow of nitrogen, heated to: 200 °C for 2 hours at 5 °C min^-1^, 285 °C at 3 °C min^-1^ for another 2 hours. Finally, the mixture was cooled to RT by removing the heat source. Under ambient conditions, an ethanol excess was added to the mixture, and a black material was precipitated and separated with a permanent magnet. The product was suspended in hexane and then precipitated with ethanol. This suspension-precipitation cycle was repeated four times, resulting in a brownish-black hexane dispersion stored at 4 °C for future uses.

### Water transfer (MF@PMAO)

The transfer into water was performed following a previously reported method with slight modifications.[33,34,67] In brief, 235 mg of poly(maleic anhydride-*alt*-1-octadecene), PMAO) were added to a flask containing chloroform (CHCl3) (195 mL). Subsequently, 5 mg of MF MNPs (in Fe content) in 5 mL CHCl3 were added dropwise, and the mixture was sonicated for 30 min. Afterwards, the solvent was slowly removed under vacuum (200 mbar, 40 °C) and 20 mL of 0.05 M NaOH were added to hydrolyze the maleic anhydride groups. The mixture was rota-evaporated (200 mbar, 70 °C) until complete evaporation of CHCl3. MF MNPs were then filtered using 0.22 μm syringe filters to remove large aggregates. To remove the excess of unbound polymer, the MNP suspension was centrifuged at 24 000 rpm for 2 h four times and finally the MNPs were redispersed in milliQ water for further use.

TAMRA labelled MNPs were obtained following the same protocol but using a PMAO polymer that was previously modified with a fluorescent dye (tetramethylrhodamine 5-(and- 6)-carboxamide – TAMRA -cadaverine, Anaspec).[33,68]

### Determination of iron concentration

The determination of iron concentration of the MF MNP suspensions was performed following a previously reported protocol to adjust iron concentration before each functionalization step.[69] In brief, 5 μL of MF MNPs were diluted in 45 μL solvent of (hexane or water) and digested with aqua regia (100 μL) solution (3:1 HCl: HNO3) at 60 °C for 15 min. Then, the samples were diluted up to 300 µL with miliQ water. At this point, 50 μL (in triplicate) was used for the iron quantification by mixing the digested samples with 60 μL of 0.25 M 1,2-dihydroxybenzene-3,5-disulfonic acid. This molecule forms a colored complex with iron, which was measured by UV/Vis spectrophotometry at λ = 480 nm using a hybrid multimodal plate reader (BioTek Synergy H1) and compared with a standard calibration curve obtained with solutions of known iron concentrations (0-1000 μg Fe mL^-1^). **Determination of antibody pI**. The pI determination was performed using a ready-to-use polyacrylamide mini gel for IEF (pH gradient 3 - 9) on the PhastSystem (Cytiva, former GE Healthcare). Abs (5 μg) were loaded and separated according to the method in Separation Technique File No. 100. After fixation with 10% trichloroacetic acid, proteins were stained with silver nitrate following manufacturer’s protocol. The calibration kit allowed pI determination in a 3–10 interval.

### Oriented and covalent conjugation of Piezo1 antibody (MF@AbPIEZ)

To perform the covalent immobilization in an oriented fashion, 4 μmol of 1-ethyl-3-(3-dimethylaminopropyl)carbodiimide hydrochloride (EDC·HCl) were mixed with 2.6 μmol of *N*-hydroxysulfosuccinimide (S-NHS) in 100 μL of 2-(*N*-morpholino)ethanesulfonic acid (MES) buffer (100 mM, pH 6.5). The mixture was incubated for 10 min at RT before the addition of MF MNPs (0.1 mg mL^-1^ of Fe). The mixture was then incubated for 30 min at RT using an orbital shaker, and the excess of reagents was removed by centrifugation (13 000 rpm, 40 min, 4°C). The MNPs were resuspended in MES 10 mM pH 5 and 3 μg of AbPIEZ (Proteintech, 15939-1-AP) were added. The mixture was incubated for 2 h at RT using an orbital shaker. To remove the unbound Ab, the mixture was centrifuged at 14 000 rpm for 1 h at 4°C. The supernatant was kept for dot-blot analysis. As a control, IgG isotype antibody (Proteintech, Ref. 30000-0-AP) was attached on the MFs surface following the same procedure (**MF@AbIgG**).

### Passivation with poly(ethylene glycol) (PEG)

0.5 mg MF MNPs (in Fe content) previously functionalized with Ab were mixed with 12.5 μmol of MeO-PEG-NH2 (750 Da, Rapp Polymere) in phosphate buffer (PB) (10 mM, pH 8) for 2 h at 37 °C. The excess of polymer was removed by centrifugation at 14 000 rpm for 45 min at 4 °C.

### Dot Blot

To quantify the amount of antibody bound to the MF MNPs we followed our previously reported work, with slight modifications.[35,49] After Ab functionalization, the supernatants (SN) (3 μL) were added to a mixed cellulose ester membrane (MCE, 0.22 μm) and left to dry at 37 °C for 30 min. The same concentration of Piezo1 Ab that was used to functionalize the MFs was utilized as the total input (100% control). Not only the SN but also the functionalized MF MNPs (MF@AbPIEZ) were deposited on the MCE membrane for subsequent analysis. MF@PMAO and MF@PMAO-PEG without Ab were used as additional controls. The membrane was blocked with TBST buffer (Tris-buffered saline (TBS 1X) with 0.1% Tween-20) containing 2.5% (w/v) of bovine serum albumin (BSA) at 37 °C for 1 h. After blocking, four washing steps were performed at 37 °C for 5 min using TBST. Then, membranes were incubated with a secondary antirabbit antibody conjugated with HRP (Dako, Ref. P0448, 5 μg mL^-1^) in TBST with 1% (w/v) BSA at room temperature for 2 h. Afterward, 4 washes with TBST (5 min each) were performed. The membrane was revealed using SuperSignal West Femto kit following the manufacturer’s protocol and the results were visualized using a Chemidoc imaging system (Bio-Rad).

**Physicochemical characterization of MF MNPs** is shown in Section 1 of the Supporting Information.

### Customized magnetic applicator

The development of the magneto-mechanical applicator was based on the NeoMag (60Nd, Madrid, Spain) device.[41] The system was originally developed to actuate on magneto-active substrates rather than actuating directly on magnetic particles attached to the cells. This conceptual change required the system adaptation and integration with the imaging tools used herein. The resulting device consists of a platform of four independently controllable sets of permanent magnets surrounding the sample, with two sets aligned along each axis (Figure 2). By controlling the relative position of these magnets with respect to the target actuation area, the resulting field over the region of interest can be modulated. This control is based on computational simulations that have been experimentally validated.

### Computational model to guide magnetic actuation

The magnetic fields generated by the magneto-mechanical applicator were simulated using COMSOL Multiphysics software, employing the Magnetic Fields, No Currents interface from the AC/DC Module. The system geometry was constructed considering NdFeB permanent magnets (grade N42H) with dimensions of 10 × 10 × 3*n* mm, where *n* represents the number of magnets in each set of permanent magnets. In addition to this configuration, an air domain was created surrounding the entire assembly to accurately model the decay of the magnetic field in the surrounding space and to properly define the boundary conditions of the simulation. The north-south orientations of the magnets were defined to match those used in the magneto-mechanical applicator, ensuring that the simulated magnetic field distribution accurately reflected the real configuration.

### Experimental characterization of the magnetic field generation and model validation

The magnetic flux density was measured over the region of interest using a gaussmeter (PCE-MFM 3000). The field components were measured along two directions coinciding with the actuation axes. These measurements were collected and compared to the numerical predictions, providing a maximum deviation of 7%.

### Cell culture

Human Umbilical Vein Endothelial Cells (HUVEC) were purchased from CLS services (Ref. 300605). Cells were maintained in Endothelial Cell Growth Medium (ECGM) (Cell Application, Ref. 211-500) at 37 °C in a humidified atmosphere containing 5% CO2. Cells between passages 2 and 6 were used for all experiments to ensure optimal characteristics and experimental consistency.

### Fluorescence microscopy

All fluorescence microscopy images were obtained on a Nikon TiE inverted microscope equipped with a sCMOS camera (Sona 4B, Andor). Appropriate excitation and emission filters were selected for each fluorophore and illumination was performed with a fluorescent lamp (C-HGFIE, Nikon). The experiments were performed using the indicated objectives; 10X-0.3NA air, 20X 0.5NA air, 40X 0.75NA air or 60X 1.4NA oil.

### Characterization of the endogenous expression of Piezo1

#### Immunofluorescence

Cells were seeded on sterile glass coverslips at a density of 5 x10^4^ cells/well in ECGM. After 24 h, cells were gently washed with PBS and incubated with Piezo1 polyclonal primary antibody AbPIEZ (1:50 Proteintech, Ref. 15939-1-AP) in ECGM for 40 min at 37 °C or with IgG antibody (1:100, Proteintech, Ref. 30000-0-AP). Afterwards, cells were washed thrice with PBS and subsequently incubated with Alexa Fluor 488 goat anti-rabbit secondary antibody (1:500, Invitrogen, Ref. A11008) diluted in ECGM (40 min at RT). Then, cells were washed again thrice with PBS and fixed with 2% PFA in PBS for 10 min at RT. Nuclei were stained with DAPI (Sigma Aldrich, Ref. 28718-90-3) diluted 1:1000 for 10 min at RT. Samples were mounted with FluorSave™ Reagent (Merck, Ref. 345789) and allowed to dry overnight. Images were obtained using the 20X objective.

#### Flow cytometry

Cells were harvested with trypsin (3 min at 37 °C), pelleted and treated with either AbPIEZ or its isotype control AbIgG (1:50 in PBS 1% BSA, Sigma-Aldrich, Ref. 9048-46-8) for 90 min at RT. Then, cells were washed with PBS and incubated with an Alexa Fluor-488 secondary antibody (Invitrogen, Ref. A11008, 1:200 in PBS 1% BSA) for 60 min at RT. For total Piezo1 expression analysis, cells were previously fixed (4% PFA, 10 min at RT), permeabilized with 0.1% of Triton X-100 for 5 min at RT, and blocked with 2% BSA and 10% goat serum in PBS for 1 h at RT. Following Ab incubations, cells were washed and resuspended in 200 μL of PBS, and their fluorescence intensity was measured on a CytoFLEX flow cytometer (Beckman Coulter), using the 488 nm laser and the FITC channel for the detection of Piezo1 (AF 488 emission at 525/40 nm). Live cells were first gated on forward scatter area (FSC-A) vs side scatter-area (SSC-A), followed by side scatter-height (SSC-H) vs SSC-A, to gate on single cells. Geometric Mean Fluorescence Intensity (MFI) was analyzed using histogram plots. Data was processed using CytExpert 2.4 and Kaluza 2.1 software.

### HUVEC labelling with MF MNPs

#### Fluorescence microscopy

HUVEC cells were seeded (5x10^4^ cells /well) on glass coverslips for 24 h in ECGM. Incubation with MFs (50 μg mL^-1^) was performed at 37 °C for 40 min in M1 medium (M1 medium containing: 20 mM HEPES, 150 mM NaCl, 5 mM KCl, 1 mM CaCl2, 1 mM MgCl2 and 2 mg mL^-1^ D-glucose, pH =7.4). MF@AbIgG or MF@PMAO-PEG were used as controls at the same concentration and incubation time. Then, cells were washed thrice with M1 medium and fixed with 4% PFA for 10 min at RT, permeabilized with 0.1% Triton X-100 for 20 min at RT and stained for 40 min at RT with Alexa Fluor 488 phalloidin (Invitrogen, Ref. A12379) diluted in 2.5 % w/w BSA in PBS. Nuclei were stained with DAPI (5 μg mL^-1^) for 10 min at RT and then the coverslips were mounted on glass microscope slides with FluorSave™ Reagent. Images were obtained using the 40X and 60X objectives.

#### Scanning Electron Microscopy (SEM)

Cells were seeded (5x10^4^ cells /well) on glass coverslips for 24 h in ECGM, washed with PBS and incubated with MF MNPs as mentioned above. Then, cells were washed several times with 0.1 M cacodylate buffer (pH 7.4) and fixed for 2h at 4°C with 0.1 M sodium cacodylate containing 8% glutaraldehyde. After 3 washes using 0.1 M cacodylate buffer (5 min), samples were dehydrated with 1 mL of methanol solutions (30, 50, 70 and 100%, 5 min each). Samples were attached on conductive double sided carbon tabs/tape and coated with a palladium layer (7 nm thickness) and imaged using a SEM microscope (INSPECT-F50) at the Advanced Microscopy Laboratories (LMA) at the University of Zaragoza.

### Calcium Imaging during magnetomechanical stimulation and analysis

For real-time calcium imaging, a 35 mm dish with a N° 1.5H glass coverslip bottom for high-quality imaging was used (ibidi, Ref. 81158). A silicone insert of 4 wells (ibidi, Ref. 80469) was attached to allow us to test 4 experimental conditions at the same time. The dish was coated with laminin (5 μg mL^-1^) in Dulbecco’s phosphate-buffered saline (DPBS), 2 h at 37 °C or overnight at 4 °C before seeding the HUVEC cells (1 x 10^4^ cells per well). After 4 h, cells were incubated with MF@AbPIEZ or MF@AbIgG (10 μg mL^-1^ in Fe content) in M1 medium for 40 min. Afterwards, 60 μL of 1 mM Fluo-4 AM loading solution (prepared following manufacturer instructions) was added (Fluo-4 Calcium Imaging Kit, Molecular Probes, F10489) for 15 min at 37 °C. Cells were washed with M1 and imaged in an M1 medium supplemented with 1 mM of probenecid (Invitrogen, F10489 D). In cases where inhibitor GsMTx-4 (Abcam, AB141871) was used, cells were pretreated for 30 min with 5 μM of GsMTx-4 in M1 medium before incubation with MFs and refreshed after washing the excess of MFs. Yoda1 (5 μM in M1 medium, MCE-HY-18723) was added 90 seconds after recording.

Images of Fluo-4 fluorescence intensity were captured using the 10X 0.3 NA objective with and additional 1.5X tube lens magnification at 480 nm excitation wavelength for 30 min at 30 second or 5 second intervals. The time-lapse images were analyzed using Fiji software after transformation and alignment using StackReg plugin (See Section 6 in the Supporting Information).

To quantitatively analyze intracellular activity based on calcium fluorescence signals, we developed an automated computational workflow using MagCalcium code. A detailed description is provided in Section 6, Supporting Information. For each experiment, more than 50 cells were considered for analysis. Calcium fluorescence intensity traces (ΔF/F0) were normalized by baseline (F0) for each cell. To normalize the signal, a lower envelope was subtracted from the ΔF/F0 traces. For somatic Ca^2+^ events detection two criteria were applied: peaks with a prominence of both > 5% ΔF/F0 and 3 standard deviations above F0. At least three independent experiments were conducted. Cumulative Ca^2+^ was measured as the finite sum of the area under the resting Ca^2+^ waveform during the stimulation period.

### Piezo1 silencing using siRNA

To assess the silencing efficiency, 15 x 10^4^ HUVEC cells were plated in a 35 mm ibidi dish in 1 mL of ECGM. After 24 h cells were washed with PBS and transfected with 1 μM of Piezo1 siRNA (Accell Human Piezo1 SMARTpool-E-020870-00-0050, Dharmacon) or NT siRNA (Accell Non-targeting Pool-D-001910-10-50, Dharmacon) using Accell siRNA Delivery Media (Dharmacon, Ref. B-005000), following manufactureŕs instructions. Piezo1 mRNA silencing was assessed at different time points after transfection (48, 72 and 96 h) using RT-qPCR. For real-time Ca^2+^ imaging, Accell delivery medium was replaced by ECGM 48 h post-transfection and cells were stimulated and imaged 72 h post-transfection.

### Viability assays after magnetomecanical stimulation

***Live/Dead assay*** was performed using the LIVE/DEAD^®^ Viability/Cytotoxicity Kit (Invitrogen, Ref. L3224). 1 x 10^4^ cells were plated on a 35 mm dish with a 4-well silicone insert in 100 µL of ECGM. 24 h after seeding, cells were treated with MF@AbPIEZ (50 µg mL^-1^ in Fe content) for 40 min and then subjected to a sustained magnetic stimulation for 40 min. After treatment, the cells were washed with DPBS, stained with 2 µM Calcein AM, and 4 µM ethidium homodimer (EthD-1) for 20 min at RT. Calcein dye is retained within live cells and produces an intense uniform green fluorescence in live cells (ex/em ∼495 nm/∼515 nm), while EthD-1 enters cells with damaged membranes producing a bright red fluorescence in dead cells (ex/em ∼495 nm/∼635 nm). After staining, the cells were washed with DPBS and immersed in (80 µL) of DPBS for fluorescence microscopy imaging. Images were obtained using a 20X objective.

#### TUNEL Assay for Apoptosis Detection

Apoptotic DNA fragmentation was assessed using the in situ Direct DNA Fragmentation (TUNEL) Assay Kit (Abcam, Ref. AB66108), following the supplier’s protocol with slight modifications. 15 x 10^4^ HUVEC cells were treated with (50 µg mL^-1^) MF@AbPIEZ alone or in combination with magnetic field for 40 min. After 24 h of treatment, cells were harvested by trypsinization, washed with cold PBS, and fixed in 1% PFA in PBS for 15 min at 4 °C. Cells were then permeabilized in ice-cold 70% ethanol for at least 30 min on ice, washed twice with the provided washing buffer and resuspended in the staining solution (FITC-dUTP, terminal deoxynucleotidyl transferase -TdT, and reaction buffer) for 1 h at 37 °C in the dark. After staining, the rinse buffer was added, and cells were centrifuged and washed twice. Fluorescence was analyzed by flow cytometry using a 488/520 nm (excitation/emission) filter set. Geometric mean fluorescence intensity was used to determine and quantify the apoptotic cells. A positive control, consisting of camptothecin-treated human lymphoma cells provided by the manufacturer, was included to validate assay performance.

#### F-actin staining and analysis of the actin content and distribution

HUVEC cells were seeded (15 x 10^3^ cells/well) on a 35 mm ibidi dish glass bottom with a 4-well silicone insert for high-quality imaging and let to spread for about 6 h before overnight serum starvation. Afterwards, cells were treated with (50 μg mL^-1^) of either MF@AbPIEZ or MF@AbIgG for 40 min in M1 medium and then subjected to magnetomechanical stimulation by placing the magnetic applicator inside a cell incubator for 20 min. Immediately after, cells were fixed with 4% PFA for 10 min at RT, permeabilized with 0.1% Triton X-100 for 20 min at RT and stained with Alexa Fluor 488 phalloidin (Invitrogen, Ref. A12379) for 40 min at RT diluted in 2.5 % w/w BSA in PBS. Nuclei were stained with DAPI (5 μg mL^-1^) for 10 min at RT. Samples were mounted with FluorSave™ Reagent on a coverslip and left to dry overnight. Images were obtained using the appropriate filter sets and using the 60X 1.4 NA oil objective.

#### Analysis

Each cell was manually outlined using a combination of phase imaging and a cytosolic marker. To estimate the F-actin distribution, each cell ROI was subdivided into a smaller central ROI and a surrounding ring-shaped ROI. The ring width, and the corresponding reduction of the central ROI, were set to 5 microns. Calculating the ratio between the ring and central ROI values allows assessment of F-actin redistribution: a ratio greater than 1 indicates enrichment toward the outer cellular cortex, while a ratio less than 1 suggests central accumulation. The actin cable intensity was measured by first identifying all the actin fibers using filament sensor. [70] After identifying the intensity distribution of the control condition, the threshold for obtaining the 5% most intense fibers was determined. Next, the fraction of fibers having intensities above the threshold were measured in distinct conditions.

#### c-Fos immunofluorescence

HUVEC cells were plated (15 x 10^3^ cells/well) into a laminin-coated 35 mm ibidi glass bottom dish with a 4-well insert, let spread for about 5 h in ECGM, and then, serum starved overnight. Afterwards, cells were treated with 50 μg mL^-1^ of either MF@AbPIEZ, MF@AbIgG or MF@PMAO-PEG for 40 min in M1 medium. The positive control (PC) was generated by treating cells with 20% FBS for 30 min at 37 °C. Sustained magnetomechanical stimulation was performed by placing the magnetic applicator inside a cell incubator for 60 min at 37 °C, 5% CO2. Immediately after treatments, cells were fixed with 4 % PFA for 10 min, permeabilized with 0.1% Triton X-100 for 20 min, and pre-blocked with PBS containing 3% bovine serum albumin and 1% normal goat serum (NGS; Vector Laboratories) for 1 h. Then, the primary antibody (anti-c-Fos, 1:250, Ref. ab190289, Abcam) was incubated for 90 min in 1% BSA in DPBS. After 2 washes with PBS, the corresponding secondary antibody Alexa Fluor 647 polyclonal donkey antibody (1:500, Abcam) were applied for 1 hour at 37 °C and counterstained with DAPI (5 μg mL^-1^) in PBS for nucleus detection for 10 min at RT. Stained cells were mounted with FluorSave™ Reagent and imaged using the appropriate fluorescence filter sets and using the 20X objective. Pearson correlation coefficient (P > 0.35) was used as a colocalization measurement using Cell Profiler program.

#### RT-qPCR

15 x 10^4^ cells were plated in a 35 mm ibidi dish and after 24 h they were subjected to a 40 min treatment with MF@AbPIEZ (10 μg mL^-1^) and 20 min of sustained field exposure. The MNPs incubation was carried out in M1 medium. At the end of the treatment, RNA was extracted using the Total RNA Purification Kit (Norgen Biotek, Ref. 17200), following the manufacturer’s instructions. The quantification of the extracted RNA was performed by measuring the absorbance at 260 nm using a Synergy H1 Microplate Reader (BioTek), while its integrity was assessed through electrophoretic run on a 1% Agarose gel in TAE buffer (40 mM Tris-Acetate; 1 mM EDTA). cDNA was synthesized using the High-Capacity cDNA Reverse Transcription Kit (Applied Biosystem, Ref. 4368814) according to the manufacturer’s instructions. RT-qPCR was conducted on four biological replicates for four experimental conditions (Control, Field, MF@AbPIEZ, MF@AbPIEZ + Field). RT-qPCR was performed using NZYSupreme qPCR Green Master Mix (NZYtech, Ref. MB419) in a CFX Opus 96 Real-Time PCR System (Bio-Rad) with the following steps: 50 °C for 2 min, 95 °C for 5 min, 95 °C for 15 seconds (denaturation); 60 °C for 1 min (annealing), repeated for 40 cycles of denaturation and annealing. Additionally, to check for the formation of undesired amplification products, a melting curve analysis was performed with a 0.5 °C increment from 60 °C to 95 °C. GAPDH was used as an internal control to evaluate the relative gene expression (Table S2, Supporting Information) using the delta–delta Ct method (2−ΔΔCT).

### Statistical analysis

All statistical analyses were performed using GraphPad Prism 8. Comparison between two groups of quantitative variables were performed using Student’s t-test. In the case of three or more groups data was analyzed one-way analysis of variance (ANOVA), followed by a multiple comparison test to compare experimental groups. Further details of the statistical analysis performed are given in each figure legend. In all cases, *p ≤ 0.0332; **p ≤ 0.0021; ***p ≤ 0.0002; ****p ≤ 0.0001.

## Supporting Information

Supporting Information is available from the author.

## Acknowledgements

This work has received funding from the European Union under the HORIZON-MSCA-2021-PF-01-01, Grant agreement No 101064735 and from European Research Council (ERC) under the European Union’s Horizon 2020 research and innovation programme (Grant agreement No. 853468), Project PID2021-122508NB-I00 funded by MICIU/AEI /10.13039/501100011033 and FEDER, UE and project CEX2023-001286-S funded by MICIU/AEI /10.13039/501100011033. TSvZ was supported by a fellowship from the Fundación General CSIC’s ComFuturo programme which had received funding from the European Union’s Horizon 2020 research and innovation programme under the Marie Skłodowska-Curie grant agreement No. 101034263. Authors acknowledge support from Gobierno de Aragón for funding the Bionanosurf (E15_23R) research group, the use of Advanced Microscopy Laboratory (Universidad de Zaragoza) for access to their instrumentation and expertise, and the use of the Servicio General de Apoyo a la Investigación-SAI, Universidad de Zaragoza and Unidad Técnica en Ingeniería de Microdispositivos. Authors also acknowledge Support from the European Research Council (ERC) under the European Union’s Horizon 2020 research and innovation programme and Horizon Europe programme, Grant -No. 101081713 (ISBIOMECH). We also thank Dr Ricardo García Salcedo for his help in the optimization of MagCalcium code.

## Data Availability Statement

The data that support the findings of this study are available from the corresponding author upon reasonable request. The code for calcium influx analysis is freely available from Zenodo ( https://doi.org/10.5281/zenodo.14692192)

## TOC entry

MagPiezo enables wireless activation of endogenous Piezo1 channels without genetic modification using 19 nm magnetic nanoparticles and low-intensity magnetic fields. It generates torque forces at the piconewton scale to trigger mechanotransduction in endothelial cells, standing as a novel platform to interrogate and manipulate Piezo1 activity in vitro, with strong potential for in vivo use.

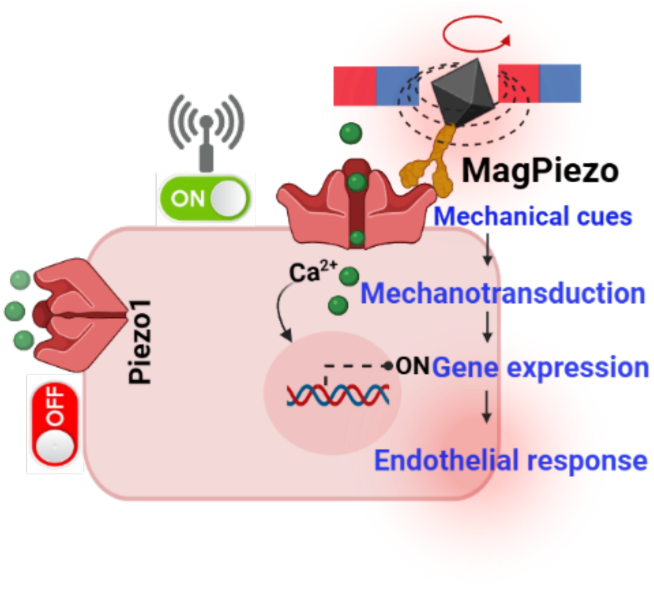

## Author Contributions

S.DS.F: conceptualization, investigation (all experiments), formal analysis, writing-original draft, review and editing, funding acquisition.

M.D.S: investigation (calcium imaging and RT-qPCR experiments), writing-review.

T.V.Z: formal analysis (F-actin analysis), methodology (real-time calcium imaging) and writing-review.

Y.F.A: methodology (simulations of magnetic applicator and theoretical calculations), writing-review and editing.

P.M.V: investigation (flow cytometry experiments), formal analysis.

D.G.G: methodology (magnetic field applicator design), writing-review and editing.

R.M.F. funding acquisition, writing-review and editing.

M.M. conceptualization, funding acquisition, supervision, writing-original draft, review and editing.

All authors discussed and commented on the manuscript. All authors have given approval to the final version of the manuscript.

