## Supporting information for "MagPiezo: a Magnetogenetic Platform for Remote Activation of Endogenous Piezo1 Channels in Endothelial Cells"

### 1. *MF MNP characterization*

The MNP shape size, and size distribution were evaluated by TEM using a Tecnai T20 (FEI, Amsterdam, Netherlands) transmission electron microscope operating at 200 kV. TEM samples were prepared by depositing 5 μl of dilute solution on a copper grid (200 mesh) followed by drying at room temperature before analysis. MNPs size distributions were obtained by measuring more than 200 MNPs using Fiji software.

High-Angle Annular Dark Field Scanning Transmission Electron Microscopy (HAADF-STEM), and energy-dispersive X-ray (EDX) images were obtained on a Tecnai F30 microscope with an accelerating voltage of 300 kV. Elemental mapping was performed using an Oxford EDS system in the drift corrector acquisition mode.

The crystal structure of the samples was identified by XRD powder patterns recorded using a Bruker D8 ADVANCE diffractometer working with CuKα ($\lambda$ = 1.5406 Å) radiation. The patterns were collected between 10° and 70° in 2θ range. Organic contents were determined by thermogravimetric analysis (TGA) using a Universal V4.5A TA Instrument (New Castle, DE, USA) under $N_2$ atmosphere at a flow rate of 50 mL $min^{-1}$ at a rate of 10 ºC $min^{-1}$ until a final temperature of 800 ºC.

The total iron and Zn, Mn and Fe content were determined by Inductively Coupled Plasma Optical Emission Spectroscopy (ICP−OES). Typically, 25 μL of MF MNPs suspension were digested in 1 mL of aqua regia in a volumetric flask at 60 °C overnight. Afterward, the flask was filled up with deionized water and samples were analyzed with an ICP plasma atomic emission spectrometer I CAP PRO XP Duo (Thermo Scientific) at the Servicio de Análisis Químico (SAI, Universidad de Zaragoza, Spain). Measurements were performed in axial mode, with the following wavelengths: Fe 259.940 nm, Mn 257.610 nm, Zn 213.856 nm. Experiments were carried out in triplicate.

Hydrodynamic diameters were obtained by Dynamic Light Scattering (DLS) using a Malvern Zetasizer Nano instrument. Samples were prepared at a concentration of 0.02 mg Fe $mL^{-1}$ in Milli-Q water and sonicated 10 s before measurement. Each sample was measured five times at 25 °C, combining 10 runs per measurement.

For the magnetic characterization, the magnetic suspensions were lyophilized and measured as powder, introduced into a gelatin capsule, and immobilized with cotton wool. Hysteresis loops were measured using a superconducting quantum interference device (SQUID, Quantum Design GmbH, Pfungstadt, Germany) magnetometer at 290 K in fields of up to 5000 kA $m^{-1}$.

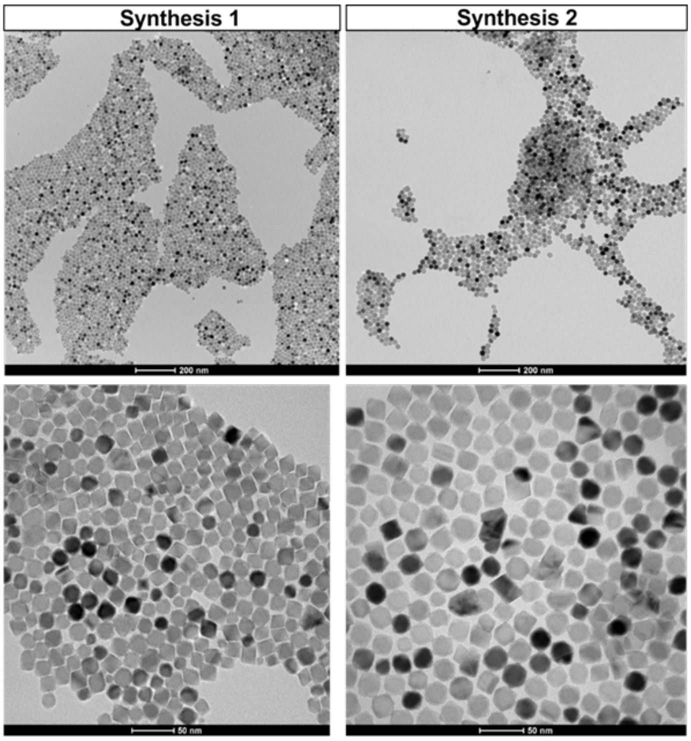


**Figure S1.** TEM images of two batches of MF@OA obtained by one step thermal decomposition method showing the reproducibility of the synthesis.

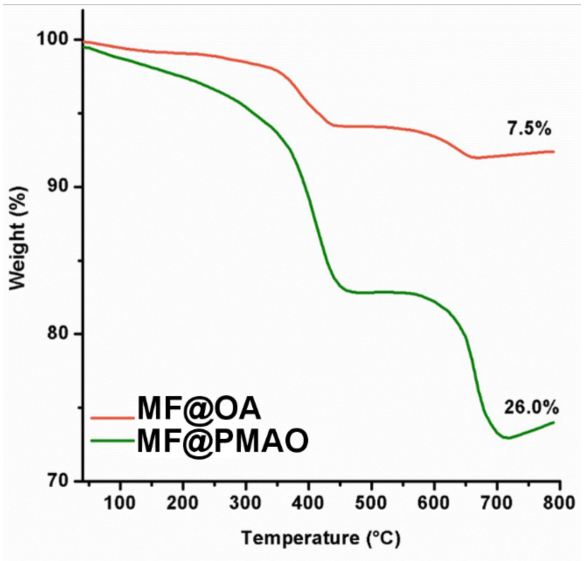


**Figure S2.** Thermogravimetric analysis (TGA) of MF@OA and MF@PMAO, respectively.

**Table S1. Magnetic properties of MF@OA and MF@PMAO measured at 300 K**

| Samples | $M_s$ ($Am^2kg^{-1}$) | $M_r$ ($Am^2kg^{-1}$) | $H_C$ (Oe) |
|---|---|---|---|
| **MF@OA** | 99.1 | 0.6 | 6.1 |
| **MF@PMAO** | 100.0 | 0.2 | 0.7 |

### *2. Characterization of the magnetic applicator*

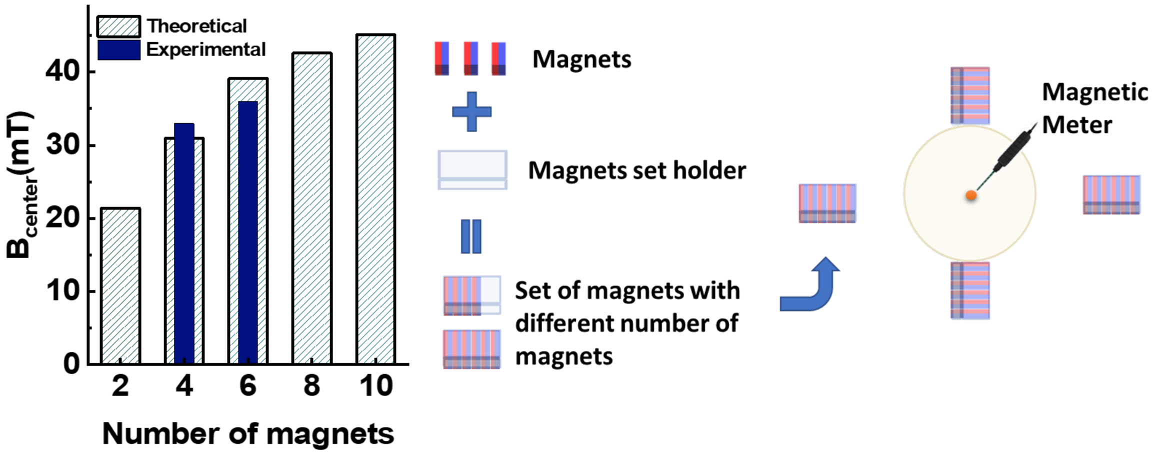


**Figure S3.** Variation of magnetic field intensity (theoretical and experimental) at the center of the petri dish as a function of the number of magnets included in each set of magnets. Numerical simulations were carried out using COMSOL Multiphysics software, considering a set of 2, 4, 6, 8 or 10 permanent magnets. Afterwards, the magnetic field intensity at the center of the circular area was experimentally measured using a gaussmeter placed at the center of a 35-mm Petri dish using sets of either 4 or 6 permanent magnets. Both experimental measurements and simulations were conducted at a fixed position (Position B) configuration of the magnetic applicator.

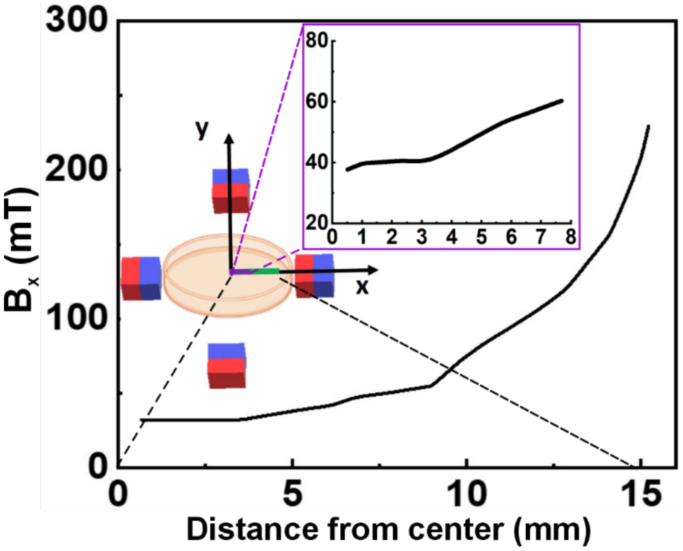


**Figure S4.** Theoretical magnetic field strength along the X axis at a fixed position (Position A configuration). The inset highlights the magnetic field intensity within the region where cells are seeded (cell culture insert of approximately 17 mm placed in the middle of the petri dish).

### 3. *Theoretical calculations of mechanical forces induced by MF MNPs*

#### *3.1. Torque. Single MF MNPs under SPM regime*

MNPs can experience different magnetic forces when they are exposed to an external magnetic field, governed by their intrinsic magnetic properties and the characteristics of the applied field. When the applied field ($\boldsymbol{B}$) is time-dependent or rotating, the particles may experience a magnetic torque ($\boldsymbol{\tau}$) that tends to align their magnetic moment ($\boldsymbol{\mu}$) with the field direction. Because the field is time-varying, the moment cannot instantaneously align. This mismatch causes a torque that tends to rotate the particle (if it's free to move) or reorient the magnetic moment within the particle. The torque $\boldsymbol{\tau}$ undergone by the MNP is calculated as:

$$\boldsymbol{\tau} = \boldsymbol{\mu} \times \boldsymbol{B} \tag{1}$$

The magnetic moment of a nanoparticle is $\mu_s = V_{\mathrm{MNP}} \cdot \rho \cdot M_s$, where $\rho$, $M_s$ and $V_{\mathrm{MNP}}$, are the density, saturation magnetization and volume of the nanoparticle, respectively.

For SPM nanoparticles, the magnetic moment fluctuates due to thermal energy, and it is not fixed. Instead, the particle exhibits a time-averaged effective moment that aligns with the external field, as described by the Langevin model:

$$\mu = \mu_s \left[\coth(\chi) - \frac{1}{\chi}\right] = \mu_s L(\chi) \;\; \text{with} \;\; \chi = \frac{\mu_s \mu_0 H}{k_B T} \tag{2}$$

where $k_B$ is the Boltzmann's constant, $\mu_0$ is the vacuum permeability, $T$ is the temperature, and $H$ is the magnetic field intensity. $L(\chi)$ accounts for the ratio between the magnetic energy of a MNP under an external field and its thermal activation.

Given our MF MNP parameters ($M_s$ = 99 $\mathrm{Am^2/kg_{ferrite}}$, $\rho$ = 5150 $\mathrm{kg/m^3}$, and $V$=3.23 X $10^{-24}$ $\mathrm{m^3}$ along with a maximum applied field $H$=36 mT (at the center of the petri dish) and a temperature $T$=37°, the dimensionless parameter $\chi$ ~14. This value indicates strong, though not complete, alignment of the magnetic moments with the external field, resulting in a small angle $\langle\theta\rangle$ between them.

To estimate the average alignment angle $\langle\theta\rangle$ we used the probability density $P(\theta)$ from the Langevin model:

$$P(\theta) = \frac{\sin\theta e^{\chi\cos\theta}}{\int_0^{\pi} \sin\theta e^{\chi\cos\theta} d\theta} \tag{3}$$

Then the average cosine of the angle is given by:

$$\langle\cos\theta\rangle = L(\chi)$$

Therefore, the average angle is:

$$\langle\theta\rangle = \cos^{-1}\big(L(\chi)\big)$$

For $\chi$ ~14, the average angle between the magnetic moment and the field is $\langle\boldsymbol{\theta}\rangle \sim \mathbf{21°}$. This misalignment is key to generating magnetic torque, as it allows the magnetic moments to experience a rotational force under the applied field.

The net torque $\tau_1$ experienced by a single MF nanoparticle is given by:

$$\tau_1 = \mu_s \cdot B \sin(21°) = 2.0 \times 10^{-20}\text{Nm} \quad \textbf{(4)}$$

The force generated by this torque is calculated as:

$$F_\tau = \tau/r \quad \textbf{(5)}$$

where $r$ (torque arm) is the position vector from the axis of rotation to the point where the force is applied. Considering the longest edge dimension of the nanoparticle (r=13.2 nm) as the torque arm, the resulting torque-induced force Equation (5) is $F_\tau = 1.5$ pN

*3.2* *Assemblies of SPM nanoparticles*

The net torque experienced by an assembly of MNPs, $\tau_{assembly}$, increases with the number of particles *N*, and is strongly influenced by magnetic anisotropy and interparticle interactions. The torque per assembly is described as:

$$\tau_{assembly} = \tau_1 N^{1-\alpha} \quad \textbf{(6)}$$

where $\tau_1$ is the torque of a single particle (See Equation (4)), and $\alpha$ is a scaling exponent that reflects the degree of magnetic alignment within the assembly. For weak dipolar interactions and random orientation, $\alpha$=0.5, resulting in $\tau_t \sim \sqrt{N}$. In contrast, partial or full alignment leads to lower $\alpha$, ($\alpha$= 0.22 or 0, respectively),[1] significantly enhancing the collective torque response as follows: $\tau_t = \tau_1 N^{0.78}$ or $\tau_t = \tau_1 N$, respectively.

Considering an assembly with an average length of 3 μm and width 0.2 μm (torque arm= 1.5 μm) which is composed by $10^4$ nanoparticles, the obtained torque is: $\tau_{assembly} = 2.6 \times 10^{-16}$ Nm at 36 mT and thus, the force due to torque generated by a single assembly is $F_{\tau_{assembly}}$= 40 pN

*4. MF MNP functionalization with antibodies*

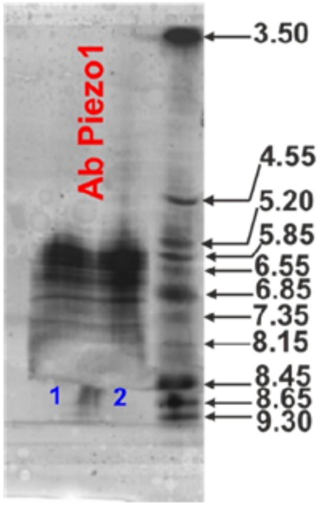


**Figure S5.** Isoelectric point (pI) of polyclonal Piezo1 antibody. Lanes 1 and 2 are technical replicates. Lane 3: isoelectric focusing marker

**Theoretical estimation of the amount of Piezo1 antibody conjugated on the MF MNPs surface**

To estimate the maximum coverage of Piezo1 antibody molecules per nanoparticle, we considered an octahedral morphology, so its volume and surface area are given by Equation (7) and (8):

$$V_{FM} = \frac{\sqrt{2}}{3} a^3 \quad (7)$$

$$A_{FM} = 2\sqrt{3} a^2 \quad (8)$$

where $a = 19 * 10^{-9}\ m$, so the volume of each MF nanoparticle and its surface area can be calculated as:

$$V_{FM} = 3.2 * 10^{-18}\ cm^3 \qquad A_{FM} = 1250.5\ nm^2$$

Typical IgG antibodies are ~ 14.5 nm x 8.5 nm x 4.0 nm in size.[2] Assuming that when using the 2-step methodology to orient the Abs they adsorb in a flat orientation,[3] the surface that they will occupy (2D projection) can be calculated as ~123.25 $nm^2$. Ideally, assuming full, flat packing and no steric hindrance of the Abs on the surface of the MF, the number of antibodies needed to fully cover the surface is of ~10 AbPIEZ antibodies per MF MNP or between 7-8 AbPIEZ considering a 70-80% surface density**.**

The mass of one MF MNPs can be calculated considering its volume and the density of $Fe_3O_4$ ($\rho$ = 5.17 g/cm$^3$): $m_{FM} = 1.67\ x\ 10^{-17} g$

Knowing the mass, the number of MNPs in 0.1 mg can be calculated as: $N_{FM} = 5.98\ x\ 10^{12}$ Finally, from the Dot Blot assay it could be concluded that all Ab added (3 μg) was attached on the MF. The number of Ab molecules in 3 μg can be obtained considering the molecular weight and the Avogadro's number ($N_A = 6.022\ 141\ 29 \times\ 10^{23} mol^{-1}$),:

$$N^{\underline{o}}\ of\ AbPiez\ molecules\ = \frac{3\times10^{-3} g}{150\ 000 \frac{g}{mol}} \times N_A = 1.22 \times 10^{13} \quad \textbf{(9)}$$

From this value it is possible to estimate the number of AbPIEZ molecules per MF MNP:

$$N^{\underline{o}}\ of\ AbPiez\ molecules\ per\ nanoparticle = \frac{1.22 \times 10^{13}}{5.98 \times 10^{12}} = 2$$

#### *5. Interaction of MNPs with HUVEC cells*

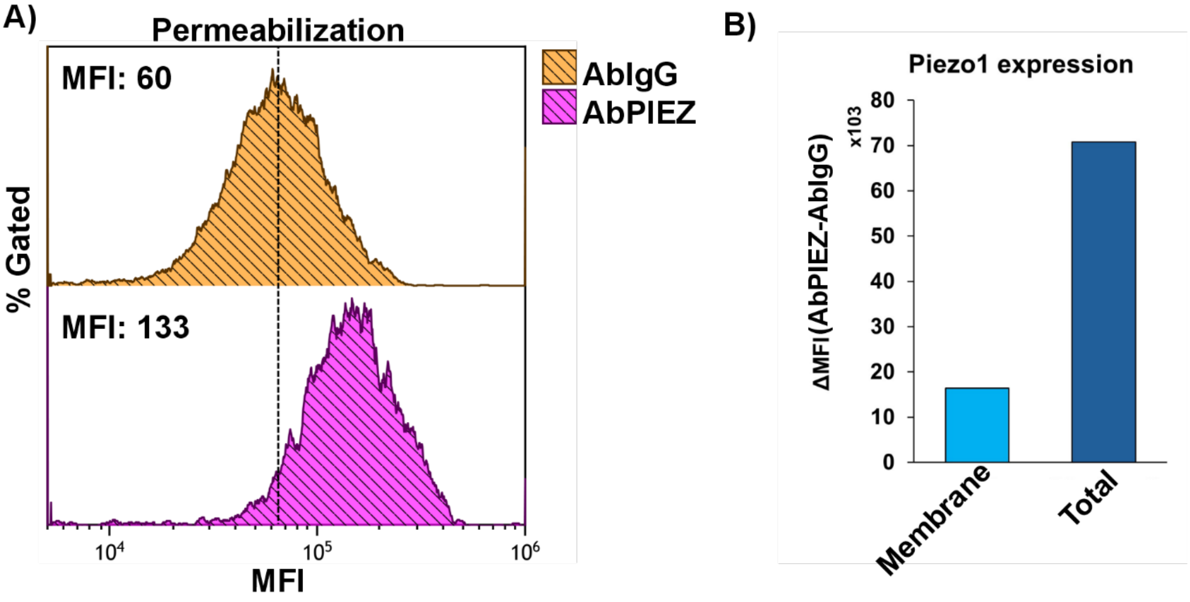


**Figure S6**. Flow cytometry analysis of Piezo1 expression. A) Flow cytometry histograms of HUVEC cells detached with trypsin, permeabilized and incubated with AbPIEZ or its isotype control (AbIgG), followed by a rabbit anti-IgG secondary antibody. Geometric Mean Fluorescence Intensities (MFI) are reported for each sample (x10³). B) Piezo1 expression levels (MFIs after subtracting isotype backgrounds) in cells analyzed under native conditions (reflecting cell membrane expression, Figure 3E) and after permeabilization (reflecting total cellular levels, Figure S6A).

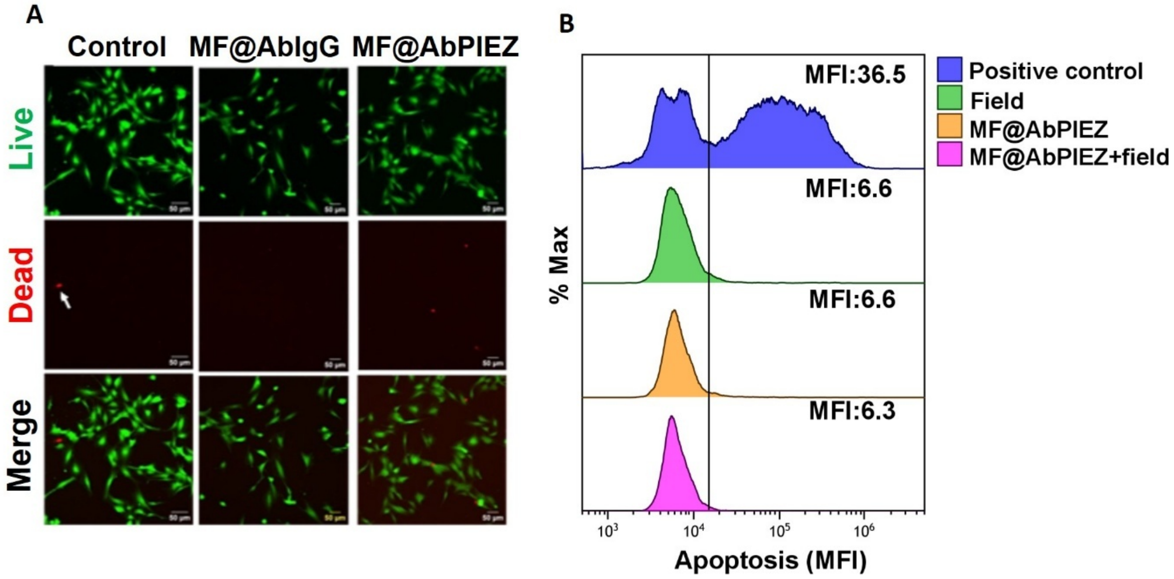


**Figure S7. A)** Representative fluorescence microscopy images of Live/Dead assay. HUVEC cells were treated with MF@AbPIEZ or MF@AbIgG for 40 minutes and exposed to sustained magnetic field for 40 minutes. Afterwards they were stained with calcein and ethidium homodimer (EthD-1) to assess the live (green) and dead (red) cells respectively. Scale bar= 50 µm. B) Apoptosis analysis by TUNEL staining using flow cytometry. HUVEC cells were treated with MF@AbPiez and exposed or not to magnetic stimulation (40 minutes). After 24 h of incubation, cells were detached with trypsin and subjected to TUNEL staining to assess apoptosis. Geometric Mean Fluorescence Intensity (MFI) values of each sample are presented $\times 10^3$.

#### *6. Calcium imaging experiments*

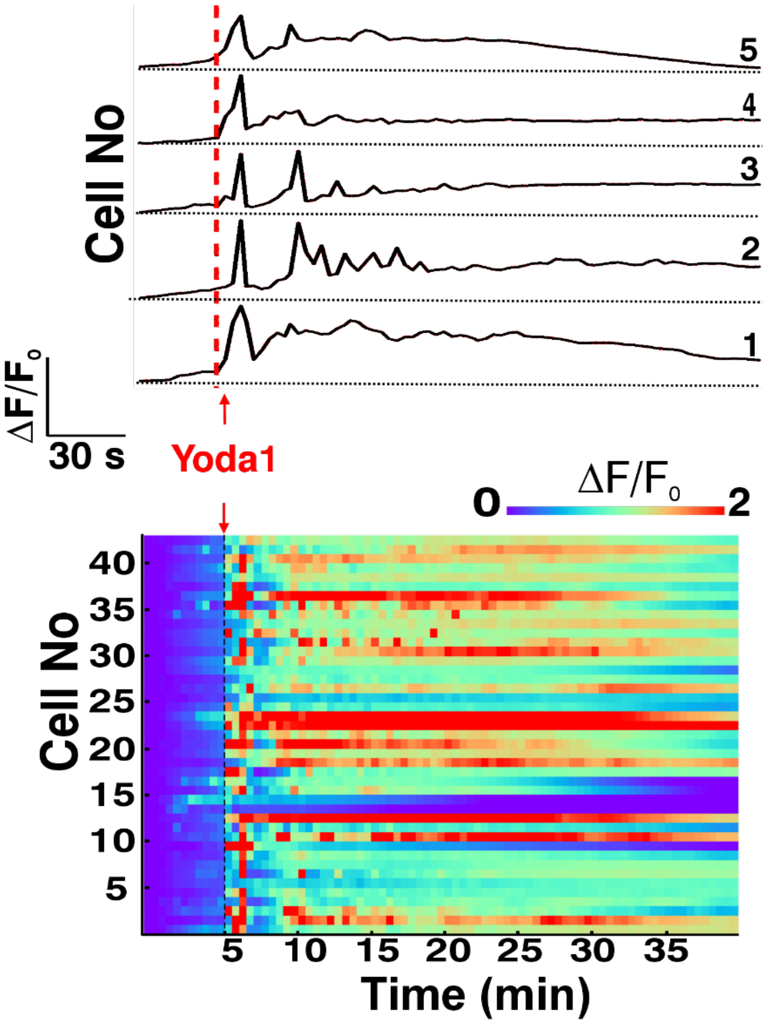


**Figure S8**. Piezo1 gating in HUVEC cellS after treatment with 5 μM of Yoda1. Upper panel: Representative normalized fluorescence intensity traces ($\Delta F/F_0$) of five cells. Lower panel: Heatmap of calcium signal (representing ΔF/F0) before and after Yoda1 addition (n = 45 cells).

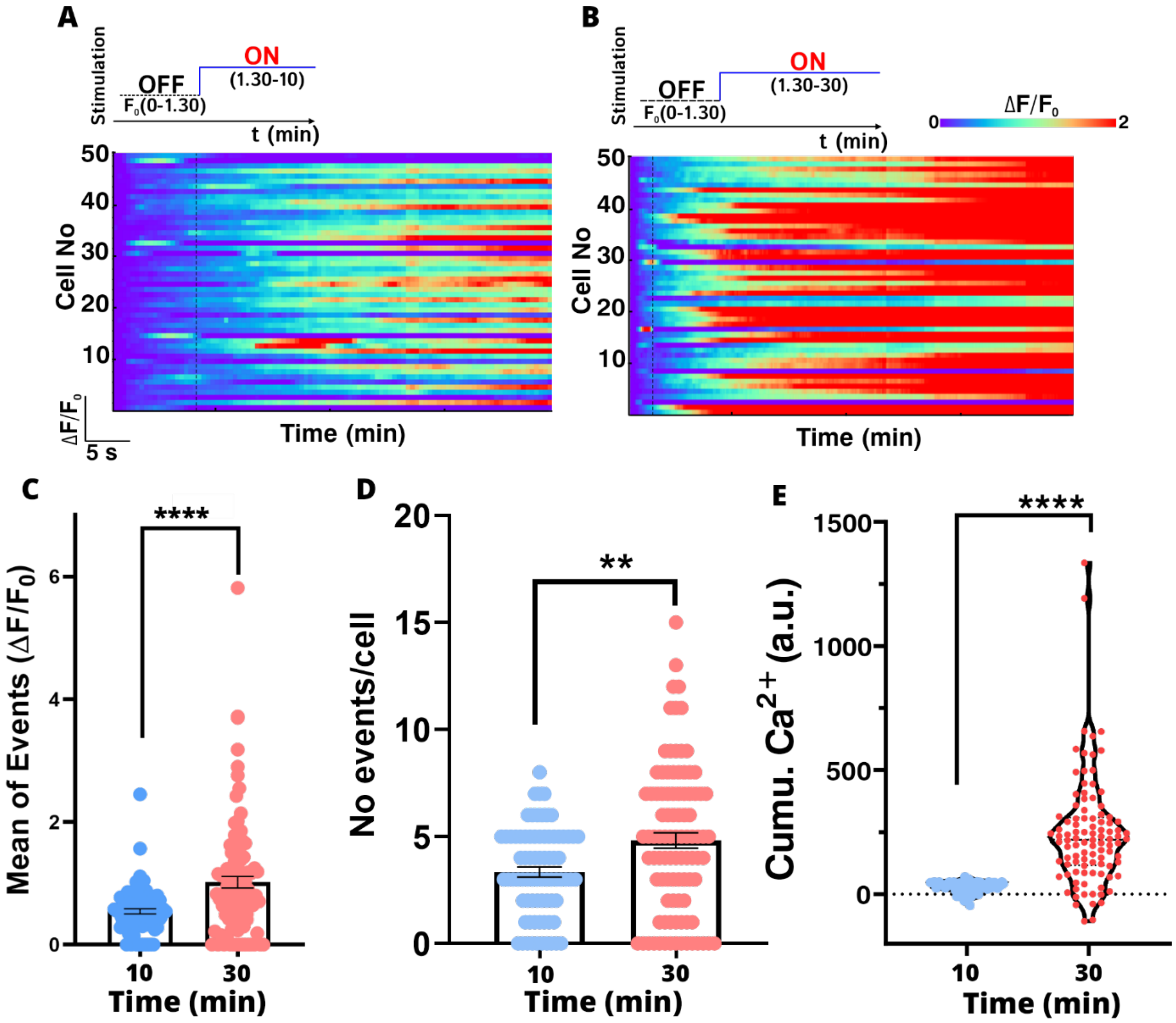


**Figure S9**. Heatmaps of HUVEC cells after A) 10 min or B) 30 min of sustained magnetomechanical stimulation. C-D) Quantification of average $Ca^{2+}$ events and number of events per cell, respectively. E) Cumulative $Ca^{2+}$ during magnetomechanical stimulation for different periods of time (10 or 30 min). Statistical analyses were performed using unpaired t-test $*p \leq 0.05$; $**p \leq 0.01$; $***p \leq 0.001$; $****p \leq 0.0001$

**Workflow for calcium fluorescence signals analysis**

To quantitatively analyze somatic calcium signals from fluorescence microscopy, we first performed motion correction followed by manual segmentation using Fiji. For signal extraction, we developed a custom computational workflow based on the MagCalcium code in Python 3.11 for downstream analysis (see Code Availability section). [4] Following MagPiezo application, we observed that the calcium traces exhibited a distinctive pattern of continuous $Ca^{2+}$ influx during magnetomechanical stimulation. [5] Therefore, we aimed to design an analysis platform that could capture this sustained signal while retaining sensitivity to discrete calcium events.

*Segmentation and Motion Correction (Fiji)*

Time-lapse fluorescence image analysis was performed in Fiji (ImageJ) using the following steps:

1. Open Raw Data and load the .nd2 file into Fiji.

2. *Spatial Binning*: Navigate to Image → Transform → Bin and apply binning with parameters: X: 4 Y: 4 Z: 1. Bin method: Sum

3. *Motion Correction*. Apply the *StackReg plugin* using the Translation transformation setting to correct XY motion artifacts. Export Preprocessed Stack and Save the motion-corrected image stack as a .tif file.

*ROI Selection and Signal Extraction*

1. Manually define regions of interest (ROIs) using the Ellipse tool within *Time Series Analyzer V3 plugin*. Select both somatic ROIs and background ROIs (from empty regions). Click More → Multi Measure to extract fluorescence intensity values over time. Cells that died or detached from the plate during the experiment were excluded from further quantification.

2. Data Export. Export the results as a .csv file containing mean fluorescence intensity values (somatic calcium traces) ($F_{raw}$) (16-bit) for each ROI across all time points.

*Data analysis (Python)*

To quantitatively analyze intracellular activity based on calcium fluorescence signals, we developed an automated computational workflow MagCalcium (See code availability). The input data, obtained from time-lapse image sequences and preprocessed with ROI segmentation in ImageJ/Fiji, were structured in Excel spreadsheets. Each file contained mean somatic raw fluorescence intensity values ($F_{raw\text{-}cell}$) and background ROIs ($F_{raw\text{-}cell}$). To obtain background-corrected fluorescence values, the script subtracts $F_{raw\text{-}cell}$ – average $F_{raw\text{-}background}$. From these corrected signals, the relative fluorescence change ($\Delta F/F_0$) was calculated as follows:

$\Delta F/F_0 = (F(t) - F_0)/F_0$,

where F(t) denotes the background-corrected signal at time t, and $F_0$ corresponds to the mean intensity over the first 6 frames (30 sec per frame interval) or 15 frames (5 sec per frame interval). $F_0$ is used as an estimate of baseline fluorescence intensity, measured during the period when magnets motion was turned off.

*Event detection*

To detect $Ca^{2+}$ events from the signals, we implemented a peak detection protocol using the findpeaks function in Python. Peaks with prominence of both >5% $\Delta F/F_0$ and 3 standard deviations of $F_0$ during the baseline period were considered events.

*Calcium accumulation*

To quantify the cumulative calcium signal for each ROI, we computed the lower envelope of the $\Delta F/F_0$ trace and performed numerical integration over time. The area under the resting $Ca^{2+}$ waveform during the magnetomechanical stimulation period was estimated using the trapezoidal method.

#### 7. *RT-qPCR experiments*

**Table S2** Set of genes assessed in the RT-qPCR study, with the corresponding NCBI accession number and primer pair used for amplification.

| Gene | Accession number | Forward | Reverse |
|---|---|---|---|
| ***VEGF-D*** | NM_004469 | GACTGGAAGCTGTGGAGATGCA | GGCTGCACTGAGTTCTTTGCCA |
| ***PCNA*** | NM_002592 | CAAGTAATGTCGATAAAGAGGAGG | GTGTCACCGTTGAAGAGAGTGG |
| ***RHOA*** | NM_001664 | TCTGTCCCAACGTGCCCATCAT | CTGCCTTCTTCAGGTTTCACCG |
| ***NFKB1*** | NM_003998 | GCAGCACTACTTCTTGACCACC | TCTGCTCCTGAGCATTGACGTC |
| ***GAPDH*** | NM_002046.7 | TGTTGCCATCAATGACCCCTT | CTCCACGACGTACTCAGCG |

#### *8. Supplementary video files.*

**Video S1** shows the custom-made magnetic applicator setup incorporated into a fluorescence microscope, as depicted in Figure 2. The device consists of 4 independent sets of 6 permanents magnets symmetrically positioned around a 35-mm circular area.

**Video S2** shows the magnetic self-assembly of MF@AbPIEZ into aligned aggregates in situ under a static magnetic field.

**Video S3** shows how MF@AbPIEZ assemblies, previously formed in the presence of a static magnetic field, tend to align their magnetic moments with the dynamic magnetic field, generating torque motion. This phenomenon is observed in various media: water, 50% v/v glycerol and on cell surfaces.

**Video S4** shows fluorescence imaging of calcium dynamics of HUVEC cells subjected to different treatments: i) control cells, ii) cells incubated with MF@AbPIEZ (AbPIEZ wo field), iii) cells exposed to the magnetic field (Field), iv) cells treated with MF@AbPIEZ and magnetostimulated (MagPiezo+) and v) cells previously blocked with GsMTx-4 and treated with MF@AbPIEZ and magnetostimulated (inh_MagPiezo+). Each frame was acquired every 30 seconds for 30 minutes and exported in .avi format at 15 frames per second. See figure 4 for analysis of the results.

**Video S5** shows fluorescence imaging of calcium dynamics of HUVEC cells treated with MF@AbIgG and exposed or not to the magnetic field. Each frame was acquired every 30 seconds for 30 minutes and exported at 15 frames per second. See figure 4 for analysis of the results.

**Video S6** shows fluorescence imaging of calcium dynamics of HUVEC cells previously treated with siRNA: i) NT siRNA; ii) Piezo1 siRNA. The videos were exported at 15 frames per second. See figure 4 for analysis of the results.
